# High-Throughput Coded Aperture Raman Imaging with Total Internal Reflection

**DOI:** 10.64898/2026.11.18.767730

**Authors:** Salvatore Sorrentino, Koseki J. Kobayashi-Kirschvink, Ke Zhang, Haochun Huang, Judy Zhou, Lloyd Bod, Jian Shu, Jeon Woong Kang, Peter T. C. So

**Author notes:** Correspondence to Jian Shu, Jeon Woong Kang, or Peter T. C. So. These authors contributed equally to this work (Salvatore Sorrentino, Koseki J. Kobayashi-Kirschvink, Ke Zhang).

## Abstract

Spontaneous Raman microscopy provides label-free molecular fingerprints of biological tissues, but its low scattering efficiency limits the throughput required for large-area hyperspectral imaging. Here we introduce total internal reflection Raman (TIRR) microscopy, an architecture that exploits repeated internal reflections within a thin sapphire substrate to generate extended evanescent excitation without dividing the incident laser power among independent illumination spots. Combined with programmable Hadamard-coded detection, TIRR recovers spatially resolved Raman spectra with an effective spatial resolution of approximately 3.5 µm and a spectral linewidth of approximately 10 cm⁻¹. Hyperspectral imaging of human skin and colon tissues reveals heterogeneous Raman contrast across submillimeter-scale fields of view. Compared with confocal Raman microscopy, TIRR achieves a directly measured 9.0-fold reduction in acquisition time at lower excitation power, corresponding to a 20.1-fold power- and signal-to-noise-normalized acquisition-efficiency advantage. We further demonstrate compressive reconstruction using low-sequency Hadamard patterns, preserving high spectral and spatial fidelity in both the development dataset and an independent tissue specimen. This approach provides an additional nominal 2.1-fold acceleration, suggesting a potential combined normalized throughput enhancement approaching 40-fold. Finally, co-registration with spatial p21 annotations in a human skin section reveals exploratory Raman spectral differences associated with cellular senescence. TIRR thus establishes a programmable approach to accelerated spontaneous Raman imaging while retaining the multidimensional spectral information needed for molecular tissue analysis.

## INTRODUCTION

Mapping the molecular heterogeneity of intact tissues requires imaging methods that retain chemical specificity over large areas and at cellular length scales^1,2^. Spontaneous Raman microscopy is particularly attractive in this context: each measured spectrum reports a molecular fingerprint without labels, with sensitivity to lipids, proteins, nucleic acids and other biochemical constituents^3,4^. Yet the weak Raman scattering process makes acquisition time the principal obstacle to scaling such measurements. At micrometer-scale spatial sampling, conventional point-scanning Raman microscopy becomes prohibitively slow for systematic tissue imaging, even when the relevant molecular contrast is readily detectable in individual spectra^5–7^.

The cost of slow acquisition extends beyond the size of an image. Because the information is distributed across many vibrational bands, a complete Raman spectrum can support analyses that go beyond a small set of chemical markers, including inference of biological states from complex spectral signatures. Recent demonstrations linking Raman measurements to gene-expression profiles, notably Raman2RNA, illustrate the value of retaining these multidimensional fingerprints rather than selecting only a few diagnostic frequencies^6,8^. Increasing imaging throughput without sacrificing the information content of spontaneous Raman spectroscopy is therefore important not only for chemical mapping, but also for its integration with emerging spatial and computational molecular profiling methods.

Parallel acquisition offers one route past the point-scanning bottleneck. Multifocus, line-scan and structured-illumination Raman microscopes interrogate several locations within the same exposure and thereby reduce the number of sequential positions that must be sampled^9–11^. Their performance, however, remains coupled to how optical power is delivered to the specimen. When a fixed excitation budget is distributed among additional focal positions, the photon flux available locally decreases; preserving Raman signal quality can then require longer integration or higher total power. Consequently, extending the illuminated region does not, by itself, guarantee a proportional gain in throughput.

Coherent Raman techniques address the weak-signal problem through a different physical mechanism and have transformed rapid chemical imaging^7,12,13^. Stimulated Raman scattering and coherent anti-Stokes Raman scattering can reach acquisition speeds far beyond conventional spontaneous Raman measurements, although the implementation of broadband or hyperspectral detection involves additional excitation and detection requirements^14,15^. Rather than attempting to compete with coherent Raman microscopy on frame rate alone, the challenge addressed here is to substantially accelerate spontaneous Raman imaging while retaining its direct spectral readout and the comparatively simple continuous-wave excitation scheme on which many Raman analysis workflows depend.

An ideal solution would simultaneously achieve the broadband spectral coverage of spontaneous Raman microscopy, the parallelized excitation of line-scan architectures, and the low-power compatibility required for near-infrared operation on biological tissues, while maintaining micrometer-scale spatial resolution. Total internal reflection (TIR) provides a natural mechanism for confining excitation to an interface while efficiently redirecting incident light. TIR Raman spectroscopy has previously been developed as a surface-sensitive spectroscopic technique, with the evanescent field providing chemically specific information from approximately the first 100–200 nm above an interface^16^. Here we present total internal reflection Raman (TIRR) microscopy, which addresses these requirements through a fundamentally different illumination geometry. In TIRR, a 785 nm diode laser is coupled into a thin polished sapphire substrate via a high-refractive-index prism. The beam propagates along the substrate by repeated total internal reflection between its upper and lower faces, generating evanescent excitation at each reflection point. Because the illumination is reused through successive reflections rather than distributed from a single pass, spatially extended excitation can be generated without proportionally increasing the incident laser power. The Raman signal is collected perpendicular to the excitation plane through a high-numerical-aperture objective, spatially separating excitation and detection before spectral filtering.

A key challenge with this approach is that the beam propagating inside the substrate has a long Rayleigh length, resulting in a lateral extent of approximately 50 µm at the sample surface—substantially larger than the spatial resolution desired for tissue imaging. To recover spatial information while preserving the throughput advantage of TIRR, we incorporate coded-aperture hyperspectral imaging using a digital micromirror device (DMD) positioned at an intermediate image plane. Sequential binary patterns encode the spatial distribution of Raman emission before it enters the imaging spectrograph. Because light originating from different DMD positions enters the dispersive spectrograph with different spatial offsets, the recorded spectrum undergoes a predictable position-dependent shift on the detector. This spatial–spectral coupling can be addressed after coded-aperture reconstruction through an internally referenced spectral correction, avoiding the need for a separately calibrated displacement map. Coded-aperture and computational Raman approaches have previously demonstrated that programmable optical encoding can increase spectrometer throughput or reduce the number of measurements required for chemical imaging^17–19^. In TIRR, we extend this concept by combining programmable spatial encoding with compressive reconstruction to explicitly trade acquisition burden against spatial and spectral reconstruction fidelity.

Here, we establish TIRR as a programmable platform for high-throughput hyperspectral Raman imaging, combining extended evanescent excitation with coded-aperture detection and computational reconstruction. We first characterize the optical architecture and demonstrate micrometer-scale spatial resolution despite an excitation profile approximately one order of magnitude wider than the reconstructed spatial features. We then extend the approach to hyperspectral imaging of human skin and colon tissues, recovering spatially heterogeneous Raman contrast and complete vibrational fingerprints across submillimeter-scale fields of view. A direct comparison with conventional confocal Raman microscopy demonstrates a 9.0-fold reduction in acquisition time at lower excitation power, corresponding to a 20.1-fold power- and signal-to-noise-normalized acquisition-efficiency advantage under the investigated conditions.

We further investigate whether acquisition time can be reduced by exploiting the spatial and spectral correlations of biological tissues through compressed sensing. By identifying the influence of Hadamard pattern sequency on reconstruction fidelity, we develop a deterministic selection strategy that reduces the measurement budget from 63 to 30 patterns while preserving high spectral and spatial similarity to the complete reconstruction. Importantly, the same pattern selection performs consistently on an independent tissue specimen, supporting its transferability across distinct biological architectures. The resulting nominal 2.1-fold reduction in integration time, combined with the experimentally derived throughput advantage of TIRR, suggests a potential normalized acceleration approaching 40-fold. Rather than relying on an assumed Hadamard multiplex signal-to-noise advantage, the proposed strategy combines parallelized excitation and detection with informed measurement selection to accelerate spontaneous Raman imaging while retaining the spectral information required for subsequent molecular analysis.

## RESULTS

### Parallelized Raman excitation by total internal reflection

High-throughput Raman imaging requires the simultaneous interrogation of many sample locations without the large reduction in local excitation that can accompany simple beam expansion. In order to accomplish this, a 785 nm beam was coupled into sapphire through a high-index prism, where successive total internal reflections generated a series of evanescent interaction sites at the specimen interface (Fig. 1A). Their spacing depends on substrate thickness, motivating a direct comparison of the observed excitation profiles.

**Figure 1.**
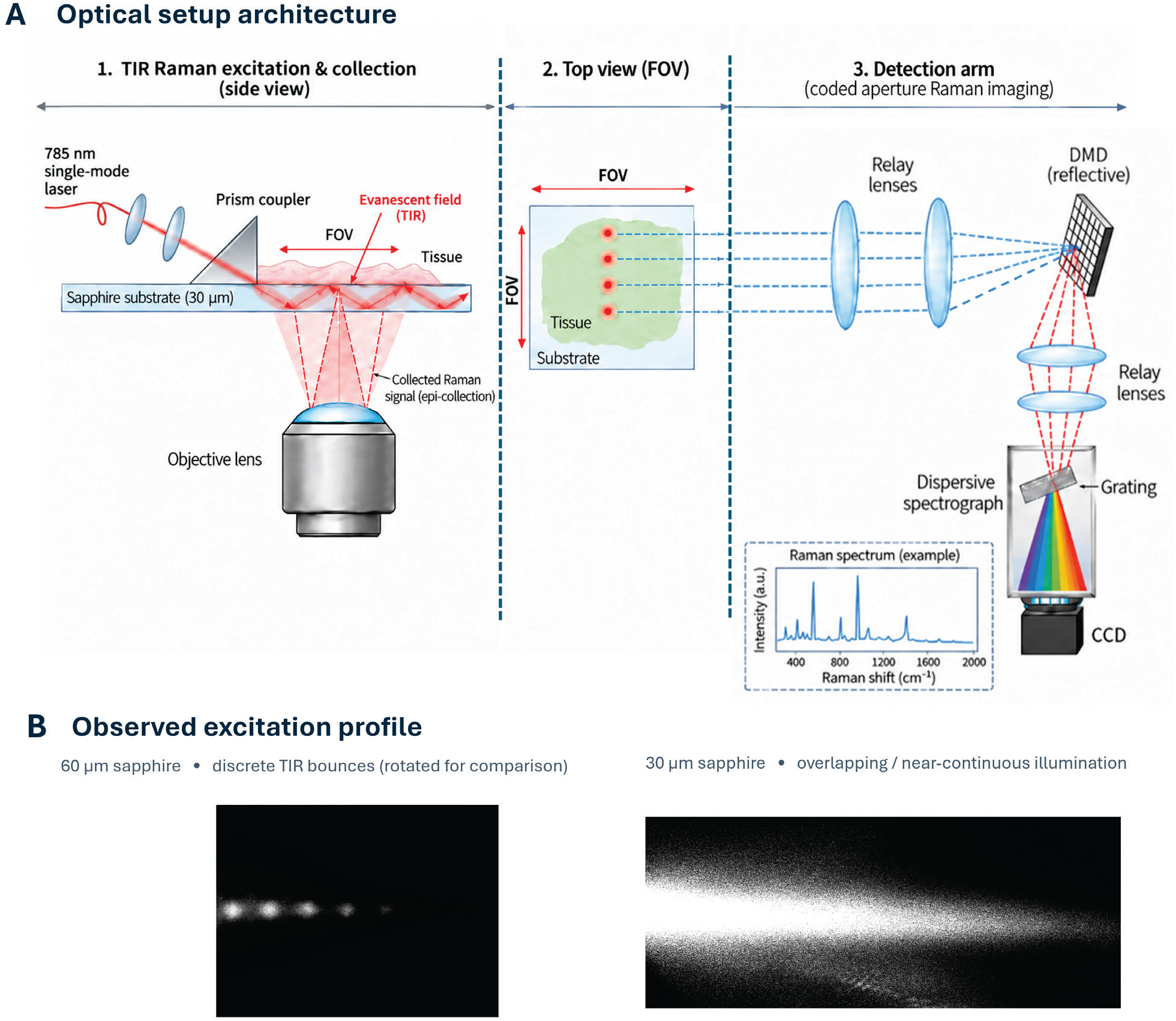
Optical architecture and extended evanescent excitation in TIRR microscopy. (A) A 785 nm beam is coupled into a 30-µm sapphire substrate and propagates by successive total internal reflections, exciting Raman scattering near the specimen–substrate interface. Raman light is collected through the substrate by a 40× objective, relayed onto a reflective digital micromirror device (DMD) for spatial encoding and then dispersed by a spectrograph onto a CCD. The top-view schematic indicates the extended excitation region within the field of view. (B) Observed excitation distributions for approximately 60- and 30-µm sapphire substrates. Reflection sites are spatially separated in the thicker substrate and overlap more strongly in the thinner substrate, yielding a more continuous excitation profile. The 60-µm image was rotated for visual comparison.

We examined this overlap by comparing the observed excitation patterns of approximately 60- and 30-µm sapphire substrates (Fig. 1B). Distinct reflection sites are visible in the thicker substrate, leaving pronounced gaps between successive excited regions. In the 30-µm substrate, their separation is reduced and neighboring contributions overlap substantially, producing a nearly continuous excitation profile across the observed region. The intensity is not perfectly uniform, but the transition from spatially isolated to overlapping excitation sites is critical: it allows the specimen to be interrogated over an extended lateral region within each acquisition rather than only at a sparse set of reflection points. We therefore adopted the 30-µm substrate for subsequent imaging experiments.

Raman emission is collected through the substrate using a 40×/0.95 NA dry objective (UPLXAPO40X, Olympus), which provides an approximately 660-µm field of view consistent with the measured value. After rejection of elastically scattered light, the image is relayed to a reflective digital micromirror device (DMD), where programmable aperture patterns encode the signal before spectral dispersion and detection on a CCD (Fig. 1A). At each position of the motorized stage, a complete sequence of coded spectral measurements is acquired simultaneously across the extended excitation region; translation perpendicular to that region supplies the second image dimension. This arrangement combines parallel excitation with spatially resolved spectral readout rather than requiring sequential illumination of every reconstructed image element.

### Coded-aperture hyperspectral imaging and spatial–spectral resolution

Maintaining an extended excitation profile over the approximately 660 µm field of view requires relatively weak focusing: tighter confinement would shorten the Gaussian beam Rayleigh length and cause rapid divergence. The resulting excitation width of approximately 50 µm is therefore considerably larger than the desired imaging resolution. To recover micrometer-scale spatial detail, we encoded Raman emission with sequential binary Hadamard masks displayed on a DMD, combining signals from multiple positions within each spectrograph exposure before computational reconstruction. Related approaches have previously been demonstrated in coded-aperture spectroscopy and hyperspectral imaging^17,20^.

In the present configuration, a complete set of 63 binary patterns was used to encode the Raman emission, with 32 of the 63 logical spatial elements contributing to each measurement. Importantly, the programmable nature of the DMD allows the spatial sampling resolution to be adjusted by modifying the size of the logical pixels and, accordingly, the number of independently encoded elements and the required Hadamard patterns. This provides a flexible trade-off between spatial sampling density and acquisition time, without requiring modifications to the optical architecture^21,22^. The spatial information was recovered by solving the corresponding linear inverse problem, as detailed in the Methods. A schematic representation of the encoding process and detection geometry is provided in Supplementary Fig. S1. An additional consideration arises from the spatial–spectral coupling introduced by the dispersive spectrograph, whereby Raman signals originating from different DMD positions undergo position-dependent spectral displacements on the detector^23^. Experimental calibration across the DMD encoding region confirmed that these displacements can be accurately approximated by a rigid spectral translation (Supplementary Fig. S2). Following coded-aperture reconstruction, individual spectra were aligned using the characteristic sapphire Raman band near 650 cm⁻¹ as an internal reference, enabling pixel-wise spectral correction without requiring a separately calibrated position-dependent displacement map.

We assessed spatial resolving performance using polystyrene microspheres with diameters of 3 and 15 µm (Fig. 2). In the mixed-bead sample, the Raman signal from an isolated 3 µm sphere remained spatially localized despite the much wider excitation profile (Fig. 2a–c). The reconstructed sampling interval was 1.15 µm per pixel, corresponding to a two-pixel sampling period of 2.3 µm. Treating the bead boundary as an effective edge under evanescent excitation, we evaluated the 10–90% intensity transition width along both reconstructed spatial axes^24^. The measured widths were approximately 3.5 µm in x and y (Fig. 2d,e), substantially smaller than the 50 µm excitation width and compatible with subcellular-scale discrimination^25^.

**Figure 2.**
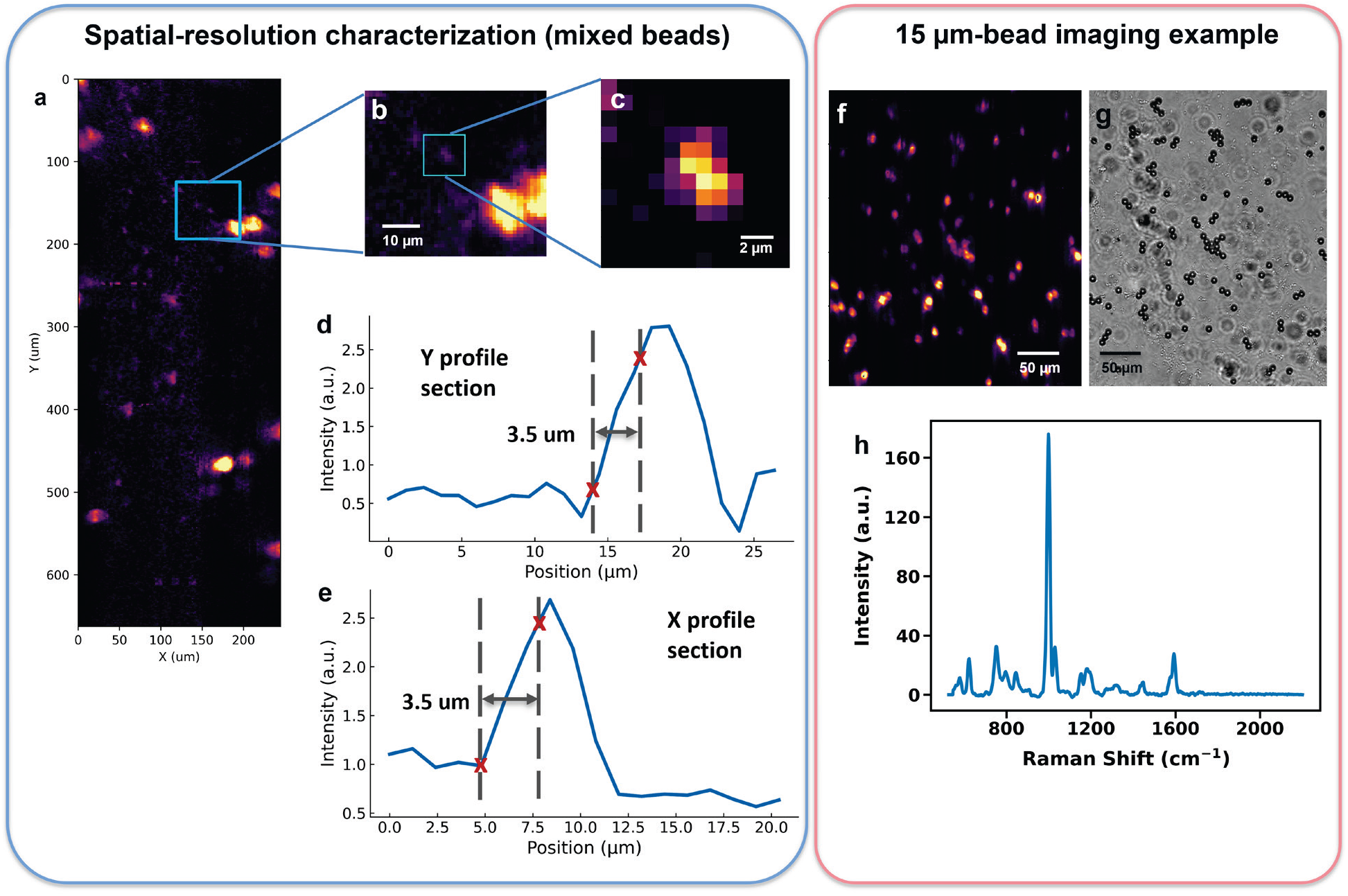
Spatial-resolution characterization and hyperspectral imaging of polystyrene microspheres using TIRR microscopy. **(a)** Reconstructed Raman image of a sample containing polystyrene microspheres with nominal diameters of 3 and 15 µm, demonstrating spatially localized Raman contrast across the imaged region. **(b,c)** Successive magnifications of the regions highlighted in (a), showing the reconstructed Raman signal from an isolated 3 µm microsphere. **(d,e)** Intensity profiles extracted along the y and x directions, respectively, across the microsphere shown in (c). The spatial resolution was evaluated using the 10–90% knife-edge criterion, yielding an effective resolution of approximately 3.5 µm in both directions, with a reconstructed pixel size of 1.15 µm. Dashed vertical lines and red crosses indicate the positions corresponding to 10% and 90% of the local intensity transition. **(f,g)** Reconstructed Raman image and corresponding brightfield micrograph of a sample containing 15 µm polystyrene microspheres, showing qualitative spatial correspondence between the Raman contrast and the bead distribution. **(h)** Representative Raman spectrum extracted from a reconstructed polystyrene microsphere, displaying characteristic vibrational bands, including the prominent peak near 1001 cm⁻¹.

To evaluate imaging performance over a larger field of view, we further examined a sample containing 15 µm polystyrene microspheres. The reconstructed Raman image resolves numerous individual beads and small clusters distributed throughout the sample (Fig. 2f). Comparison with the corresponding brightfield image shows close qualitative correspondence between the spatial distribution of the Raman features and the observed bead morphology (Fig. 2g). Beyond spatial localization, the reconstructed data retain the characteristic vibrational fingerprint of polystyrene, including the prominent band near 1001 cm⁻¹ and additional spectral features throughout the fingerprint region (Fig. 2h). These results demonstrate the recovery of both spatially resolved contrast and broadband Raman information from the multiplexed measurements.

To quantitatively assess the spectral resolving performance of TIRR, we characterized the spectral response of the detection system using narrow emission lines from a neon–argon (Ne–Ar) calibration lamp. This approach provides a direct experimental measure of spectral linewidth and has been employed in high-resolution Raman instrumentation^26^. We compared measurements obtained with the conventional spectrograph slit closed and through the coded-aperture configuration, thereby evaluating the influence of multiplexed detection on spectral resolution. Four relatively isolated emission lines, distributed between approximately 640 and 1780 cm⁻¹ on the Raman-shift-equivalent calibration axis, were selected for quantitative analysis (Supplementary Fig. S3).

For each spectral feature, the full width at half maximum (FWHM) was determined directly from the detector samples using a local linear baseline and the two pixels closest to half the peak height. The same procedure was applied to both configurations, with spectral widths expressed in cm⁻¹ using the experimental calibration (Supplementary Fig. S3 and Supplementary Note 1).

With the conventional slit closed, the four selected emission lines exhibited FWHM values between 5.8 and 8.2 cm⁻¹, yielding a mean of 7.0 ± 1.3 cm⁻¹. Under coded-aperture detection, the corresponding features showed widths between 8.0 and 11.7 cm⁻¹, with a mean FWHM of 10.0 ± 1.5 cm⁻¹ (mean ± standard deviation across four spectral lines). Taken together, these measurements establish an effective spectral resolution of approximately 10 cm⁻¹ at the investigated spectral positions under coded-aperture acquisition, compared with approximately 7 cm⁻¹ for the conventional slit-closed configuration.

The spatial and spectral characterizations thus demonstrate that TIRR combines an effective spatial resolution of approximately 3.5 µm with a coded-aperture spectral linewidth of approximately 10 cm⁻¹ at the investigated calibration positions. Together with the broadband polystyrene spectra and the spatial correspondence with brightfield imaging, these results establish the resolving capability of the platform and provide the experimental basis for evaluating its imaging throughput and subsequent application to biological tissues.

### Hyperspectral Raman imaging of biological tissues

Having characterized the spatial and spectral performance of TIRR using reference samples, we next investigated its ability to recover spatially resolved molecular information from biological tissues with heterogeneous composition and complex microarchitecture. Hyperspectral Raman measurements were performed on skin and colon tissue sections, extending the evaluation of the system from isolated molecular targets to specimens containing continuously distributed and overlapping biochemical constituents (Fig. 3). These experiments were designed to assess whether the combination of extended evanescent excitation and coded-aperture detection could preserve recognizable tissue organization while recovering local Raman fingerprints across submillimeter-scale fields of view.

**Figure 3.**
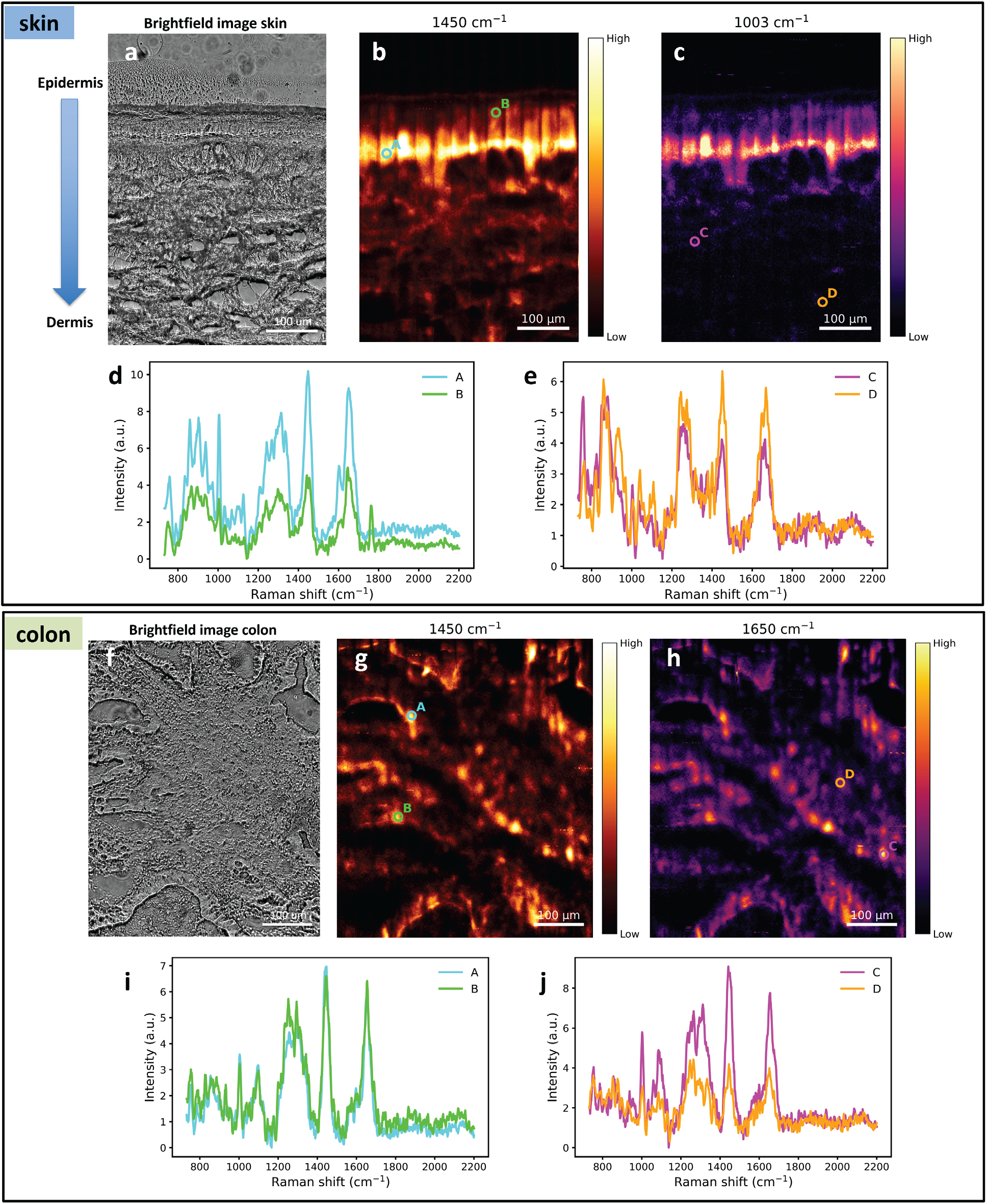
Hyperspectral Raman imaging of skin and colon tissue using TIRR microscopy. (a–e) Imaging of a skin tissue section encompassing the epidermis and underlying dermal region over a field of view of 467 × 660 µm² (391 × 552 reconstructed pixels). (a) Corresponding brightfield micrograph showing the layered tissue architecture, with the epidermal region located above the underlying dermis. (b,c) TIRR Raman intensity maps reconstructed at 1450 cm⁻¹ (CH₂/CH₃ deformation modes of proteins and lipids) and 1003 cm⁻¹ (primarily phenylalanine ring breathing), respectively. Both maps reveal a pronounced region of enhanced Raman intensity near the presumed dermal–epidermal transition. (d,e) Representative Raman spectra extracted from the spatial locations marked A–B in (b) and C–D in (c), respectively, illustrating differences in local Raman signal intensity and spectral features across the tissue. (f–j) Hyperspectral imaging of a colon tissue section over a field of view of 554 × 660 µm² (463 × 552 reconstructed pixels). (f) Corresponding brightfield micrograph showing the heterogeneous tissue morphology. (g,h) Raman intensity maps reconstructed at 1450 cm⁻¹ (CH₂/CH₃ deformation modes) and 1650 cm⁻¹ (primarily amide I vibrations with possible contributions from unsaturated lipid C=C stretching), revealing spatially distributed molecular contrast across the specimen. (i,j) Representative Raman spectra extracted from positions A–B in (g) and C–D in (h), respectively, displaying characteristic vibrational bands and local intensity variations. Both specimens were imaged using 63 binary Hadamard patterns with an integration time of 80 s per pattern and a coupled laser power of 118 mW. The datasets were acquired using nominally 50-µm-wide tiles with a 10 µm overlap between adjacent tiles. The skin dataset comprised 12 acquisition tiles and required approximately 18 h, whereas the colon dataset comprised 15 tiles and required approximately 22.5 h. Colors represent relative Raman band intensities, with independently displayed intensity scales for each map. Scale bars, 100 µm.

We first examined a skin section containing the epidermis and underlying dermal tissue over an area of 467 × 660 µm² (Fig. 3a–e). Raman images reconstructed at 1450 and 1003 cm⁻¹ revealed a pronounced spatial organization, with a particularly intense, laterally extended band near the transition between the epidermal and dermal compartments (Fig. 3b,c). The 1450 cm⁻¹ band predominantly reflects CH₂/CH₃ deformation modes associated with proteins and lipids, whereas the feature near 1003 cm⁻¹ is primarily attributed to phenylalanine residues in proteins. Interestingly, both vibrational bands exhibit enhanced intensity along a common tissue region, suggesting that the observed contrast reflects a localized change in the molecular environment rather than the spatial distribution of an isolated chemical constituent.

The position and morphology of this Raman-rich region are consistent with the dermal–epidermal junction, where basal keratinocytes interface with the specialized extracellular matrix separating the epidermis from the underlying dermis. Previous Raman imaging studies of human skin have identified distinct spectral signatures associated with the epidermis, basal epidermal compartment and dermis^27,28^. Furthermore, Raman characterization of type I and IV collagens has revealed spectral differences between the dermis, epidermis and their interface^29^. In this context, the enhanced Raman response observed here may reflect the transition between the keratin-rich basal epidermal compartment and the collagen-containing extracellular matrix of the superficial dermis. Although these contributions cannot be uniquely resolved only from the selected intensity maps, the spatial localization of the enhanced signal is consistent with the known biochemical heterogeneity of the junctional region. Representative spectra from superficial and deeper tissue locations further reveal differences in Raman signal intensity and relative spectral features (Fig. 3d,e), illustrating the recovery of local vibrational information across the layered skin architecture.

To examine the applicability of TIRR to a different tissue organization, we subsequently acquired a hyperspectral image of a colon section covering 554 × 660 µm² (Fig. 3f–j). Unlike the layered architecture of skin, the colon specimen exhibited a more heterogeneous spatial arrangement, with irregular tissue structures visible in the corresponding brightfield image. Raman maps reconstructed at 1450 and 1650 cm⁻¹ revealed spatially varying intensities that broadly followed the tissue morphology, highlighting both extended structures and localized regions of enhanced signal. The selected bands probe different, partially overlapping molecular contributions: CH₂/CH₃ deformation modes near 1450 cm⁻¹ and predominantly protein amide I vibrations, with additional contributions from unsaturated lipids, near 1650 cm⁻¹. Representative spectra acquired from four spatially separated locations retained multiple characteristic Raman features across the investigated spectral interval (Fig. 3i,j), with variations in band intensities reflecting the heterogeneous Raman response of the colon specimen. Additional TIRR acquisitions from a separate colon tissue specimen further demonstrated spatially heterogeneous Raman contrast at 1003 and 1450 cm⁻¹, together with the recovery of characteristic vibrational fingerprints (Supplementary Fig. S4).

The 12-tile skin and 15-tile colon acquisitions required approximately 18 and 22.5 h, respectively, yielding densely sampled hyperspectral datasets containing approximately 2.2 × 10⁵ and 2.6 × 10⁵ reconstructed spatial spectra. Taken together, the skin and colon measurements show that TIRR can extend programmable Raman imaging to structurally distinct biological tissues while retaining both spatially organized molecular contrast and local spectral information. This capability provides the experimental basis for evaluating acquisition throughput against conventional confocal Raman microscopy and for subsequent molecular analyses of biological tissue states.

### Raman imaging throughput benchmark against confocal microscopy

Having demonstrated hyperspectral Raman imaging across structurally distinct biological tissues, we next quantified the acquisition efficiency of TIRR relative to conventional confocal Raman microscopy. Because the two approaches differ fundamentally in their excitation and detection geometries, the comparison was designed to evaluate spectral acquisition efficiency under experimentally accessible imaging conditions rather than establish equivalent spatial or axial resolution^30^. We acquired hyperspectral images of the same colon tissue specimen over a corresponding tissue region using both TIRR and confocal Raman microscopy. The resulting datasets exhibited similar large-scale morphological features, although differences in fine spatial contrast were apparent, consistent with the distinct imaging geometries and effective sampling characteristics of the two systems (Supplementary Fig. S5).

To quantify the quality of the recovered Raman information, we compared the spectral signal-to-noise ratio (SNR) associated with the phenylalanine band near 1003 cm⁻¹, selected as a relatively narrow and well-defined vibrational feature present in both measurements. For each spatial spectrum, the Raman signal was estimated from the mean intensity within an approximately 10 cm⁻¹ interval surrounding the band, while the noise level was estimated from the spectral standard deviation in a peak-free region between 1847 and 1870 cm⁻¹. The analysis was restricted to tissue-containing pixels, excluding the surrounding substrate and low-signal regions. Representative spectra from the two modalities retained the principal Raman features throughout the fingerprint region, with differences in relative band intensities reflecting the distinct sampling conditions (Fig. 4a). The median spectral SNR was 11.0 for TIRR and 13.2 for confocal microscopy (Fig. 4b), indicating that the two acquisitions provided Raman spectra of comparable signal quality according to this operational metric.

**Figure 4.**
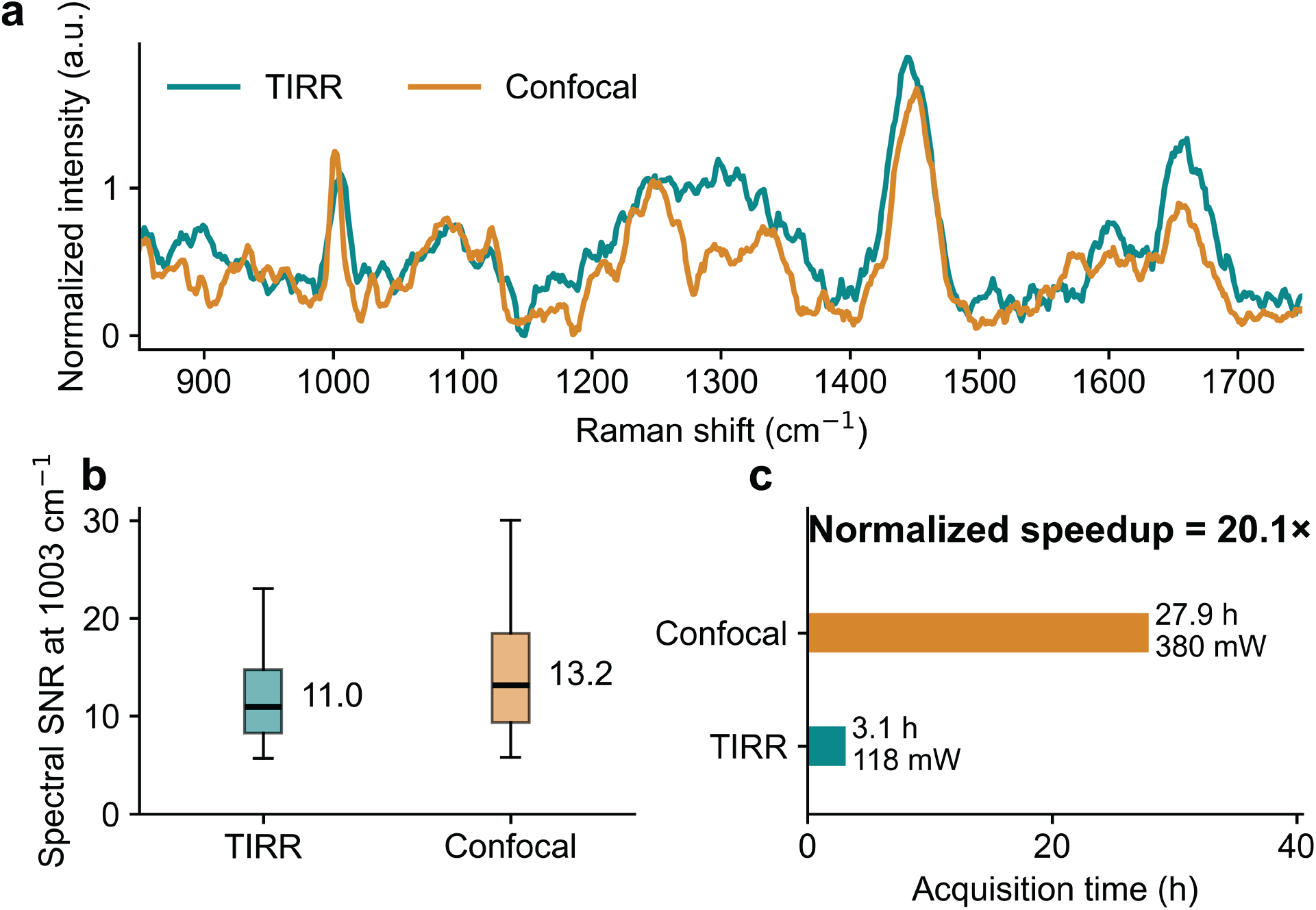
Raman imaging throughput comparison between TIRR and confocal microscopy. (a) Representative Raman spectra acquired from corresponding regions of the same human colon tissue specimen using TIRR (teal) and conventional confocal Raman microscopy (orange). Spectra were individually baseline-offset and normalized to the mean intensity of the phenylalanine band near 1003 cm⁻¹ for visual comparison. (b) Distributions of pixel-wise spectral signal-to-noise ratios (SNRs), calculated from the mean phenylalanine band intensity relative to the spectral standard deviation within a peak-free region at 1847–1870 cm⁻¹. Tissue-containing pixels were selected independently in each image to exclude substrate and low-signal regions. Boxes indicate the interquartile range, horizontal lines denote the median, and whiskers extend from the 5th to the 95th percentile. Median SNR values were 11.0 for TIRR and 13.2 for confocal microscopy. (c) Comparison of acquisition times and excitation powers for the two measurements. TIRR required 3.1 h at 118 mW, whereas confocal microscopy required 27.9 h at 380 mW. The resulting power-and SNR-normalized imaging speedup was 20.1-fold, calculated from the ratio of squared spectral SNR per unit excitation power and acquisition time. The normalized speedup is derived from experimental measurements under the stated scaling assumptions and does not imply equivalent spatial resolution or axial sampling depth.

The TIRR acquisition required 3.1 h at an excitation power of 118 mW, compared with 27.9 h at 380 mW for the confocal measurement. This corresponds to a directly observed 9.0-fold reduction in acquisition time despite the lower excitation power used by TIRR. To account for differences in both power and spectral SNR, we additionally calculated a normalized acquisition-efficiency factor based on the experimentally determined SNR, laser power and acquisition duration, whose mathematical formulation can be found in the *Methods* section. Under the conventional scaling assumption that squared SNR increases proportionally with excitation power and integration time, this comparison yielded a power-and SNR-normalized imaging speedup of 20.1-fold for TIRR (Fig. 4c). The normalized factor represents an experimentally derived estimate of the relative spectral acquisition efficiency rather than a direct measurement under identical optical conditions. Together, these results demonstrate a substantial throughput advantage for TIRR in the investigated tissue-imaging configuration, providing the basis for evaluating whether the acquisition burden can be reduced further through compressive coded-aperture reconstruction.

### Compressive hyperspectral Raman imaging through low-sequency Hadamard encoding

Following the experimental characterization of the throughput advantage of TIRR over conventional confocal Raman microscopy, we investigated whether acquisition time could be further reduced by reconstructing hyperspectral images from an incomplete set of Hadamard measurements^31,32^. Although the complete coded-aperture reconstruction uses 63 patterns, biological tissues exhibit spatial and spectral correlations that may be exploited to recover Raman information from fewer measurements^33^. To investigate this possibility, we implemented a compressed sensing reconstruction incorporating joint spatial and spectral total variation regularization, systematically evaluating the trade-off between measurement number and reconstruction fidelity. Subsets containing 10, 20, 30, 40, 50 and 60 patterns were extracted from a complete colon tissue acquisition, with ten randomly generated subsets evaluated at each sampling level. Reconstruction quality was assessed against the full-pattern reference using the spectral angle mapper (SAM) and structural similarity index (SSIM), quantifying spectral and spatial fidelity, respectively (Supplementary Fig. S6). Both metrics improved progressively with increasing measurement number, without revealing a distinct optimal sampling threshold. We selected 30 patterns as a practical operating point, reducing the measurement budget by more than 50% while retaining substantially better reconstruction quality than the more aggressively undersampled configurations.

Interestingly, reconstruction fidelity depended not only on the number of acquired patterns but also on their composition. We therefore characterized each Hadamard mask by its sequency, defined as the number of binary transitions between adjacent encoded elements. Analysis of the randomly selected subsets revealed that enrichment in low-sequency patterns was associated with improved reconstruction quality, particularly within the intermediate undersampling regime (Supplementary Figs. S7–S8). At 30 patterns, low-sequency enrichment exhibited Spearman correlations of approximately −0.47 with SAM and +0.60 with SSIM, with similarly consistent associations at 40 patterns (ρ ≈ −0.45 and +0.58, respectively). These relationships became less pronounced as the number of acquired patterns approached the complete measurement set, suggesting that mask composition plays a more consequential role when the available measurements are substantially reduced. Although the correlations across all sampling levels were weaker, the consistent behavior observed at 30 and 40 patterns provided a rationale for prioritizing slowly varying Hadamard patterns in the compressed acquisition. We therefore constructed a deterministic 30-pattern subset by selecting the masks with the lowest sequency directly from the encoding matrix, independently of their measured reconstruction performance.

This informed selection substantially improved reconstruction fidelity in the colon development dataset (Fig. 5a–d). The 30-pattern reconstruction achieved a median SAM of 7.37° and an SSIM of 0.8633 relative to the complete 63-pattern reference, compared with median values of 14.89° and 0.6865, respectively, across the ten randomly selected 30-pattern subsets. The low-sequency selection outperformed all ten random subsets in both metrics. Raman intensity maps reconstructed near 1450 cm⁻¹ preserved the main morphological features of the full-pattern image, including localized regions of enhanced signal and extended tissue structures (Fig. 5a,b). Representative spectra extracted from corresponding locations showed close agreement in the positions and relative intensities of the principal Raman bands (Fig. 5c,d), indicating that the reduced measurement set retained substantial spectral information alongside the spatial organization of the tissue.

**Figure 5.**
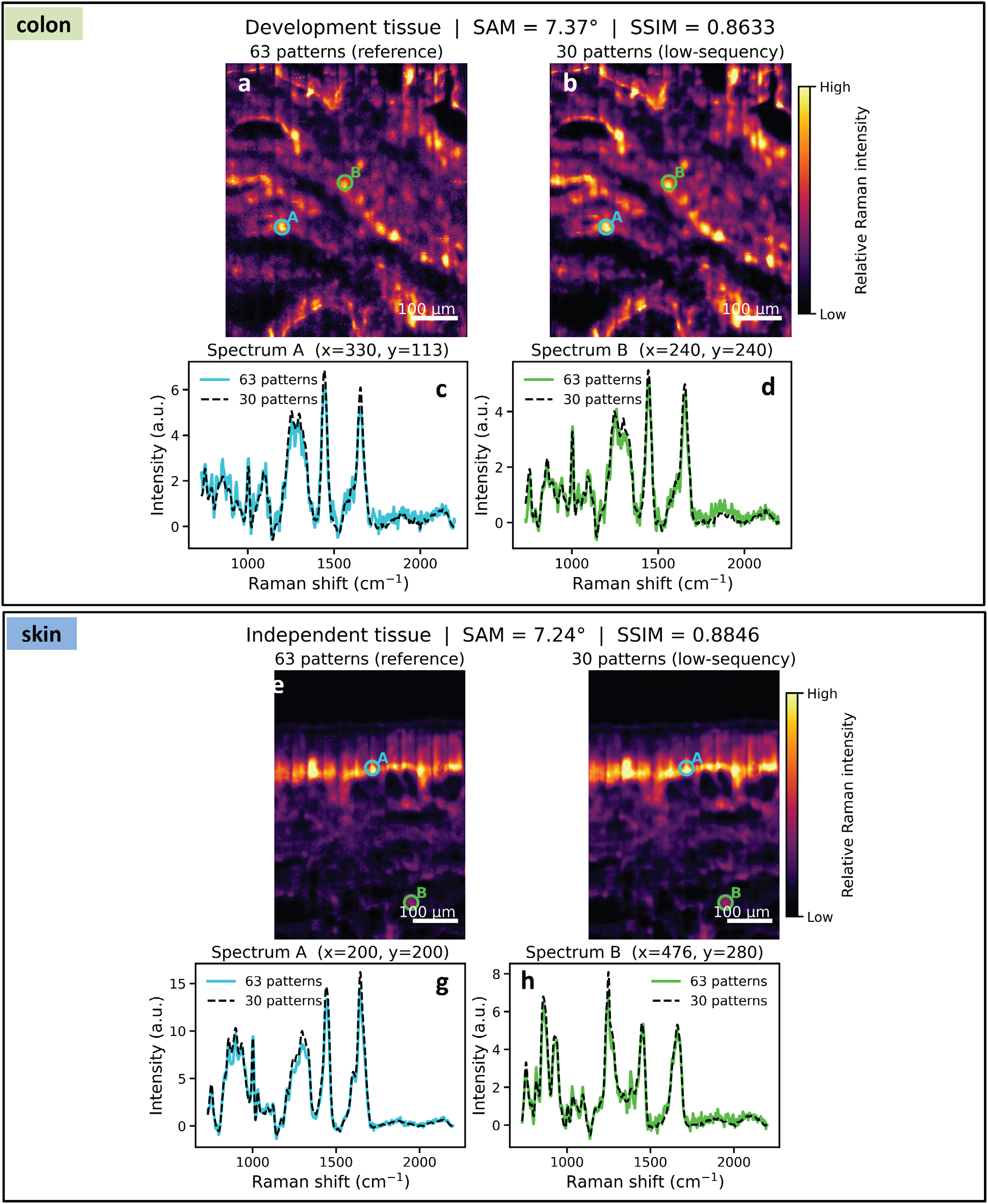
Compressive hyperspectral Raman imaging using low-sequency Hadamard pattern selection. Comparison of full-pattern (63 measurements) and compressed (30 measurements) TIRR reconstructions of human colon and skin tissues. (a,b) Raman intensity maps near 1450 cm⁻¹ reconstructed from the complete set of 63 Hadamard patterns (a) and a deterministically selected subset of 30 low-sequency patterns (b) in the colon development dataset. The reduced-pattern reconstruction preserves the principal spatial features of the reference image, achieving a median spectral angle mapper (SAM) of 7.37° and a structural similarity index (SSIM) of 0.8633. (c,d) Representative Raman spectra extracted from the corresponding positions A and B marked in (a,b), comparing full-pattern (solid colored lines) and compressed (black dashed lines) reconstructions. (e–h) Independent evaluation of the same 30-pattern subset on a skin tissue section, without modification of the mask selection or reconstruction parameters. (e,f) Raman intensity maps near 1450 cm⁻¹ reconstructed using 63 (e) and 30 (f) patterns, showing preservation of the layered tissue organization. The compressed reconstruction achieves a median SAM of 7.24° and an SSIM of 0.8846 relative to the full-pattern reference. (g,h) Representative Raman spectra from positions A and B in the skin tissue, showing agreement in the principal vibrational features between the complete and reduced-pattern reconstructions. The 30-pattern subset was selected based on the lowest sequency values of the Hadamard encoding matrix, corresponding to 47.6% of the complete measurement set. Spectral fidelity was evaluated using the median SAM over the reconstructed Raman spectra, whereas spatial fidelity was quantified using SSIM on spectral-norm intensity maps. Intensity scales are shared between the full and compressed images within each tissue dataset. Scale bars, 100 µm.

To examine whether this selection strategy was specific to the colon specimen, the same 30 Hadamard patterns were applied to an independent skin tissue dataset, without modifying the mask selection or reconstruction parameters. The resulting hyperspectral reconstruction again showed close correspondence with its complete 63-pattern reference, achieving a median SAM of 7.24° and an SSIM of 0.8846 (Fig. 5e–h). The prominent layered Raman contrast was preserved, while representative spectra from spatially distinct regions retained the principal vibrational features and their relative intensity distributions. These results support the transferability of the selected measurement basis between the two tissue architectures investigated here. Reducing the acquisition from 63 to 30 patterns corresponds to 47.6% of the original measurement budget, or a potential 2.1-fold reduction in integration time at fixed exposure per pattern. Although evaluated retrospectively using complete acquisitions, this result establishes a practical route toward further increasing TIRR imaging throughput through informed pattern selection, with a quantifiable compromise in spatial and spectral reconstruction fidelity.

### Spatially resolved Raman signatures associated with cellular senescence

To investigate the potential of TIRR for characterizing biologically defined cellular states beyond conventional chemical tissue imaging, we integrated hyperspectral Raman measurements with spatially resolved molecular annotations of cellular senescence in human skin. Following TIRR acquisition, the same tissue section was processed using STARmap-based in situ hybridization and DAPI staining to identify p21-expressing cells^34^. Spatial registration between the fluorescence and TIRR datasets enabled molecular annotations to be associated with corresponding Raman sampling regions (Fig. 6a,b). The Raman intensity map reconstructed near 1003 cm⁻¹, primarily associated with phenylalanine, revealed tissue organization consistent with the registered fluorescence image, particularly along the cellularly dense superficial region. By transferring fluorescence-derived cell segmentations into the Raman coordinate system, we extracted cell-associated spectra from 22 p21+ and 965 p21− regions. Spectra were normalized using their L1 norm across the 600–1800 cm⁻¹ fingerprint region to reduce variations in overall Raman signal intensity while preserving relative differences in their vibrational features. This multimodal strategy provided a means of investigating whether the molecular heterogeneity captured by TIRR could be related to independently identified cellular phenotypes.

**Figure 6.**
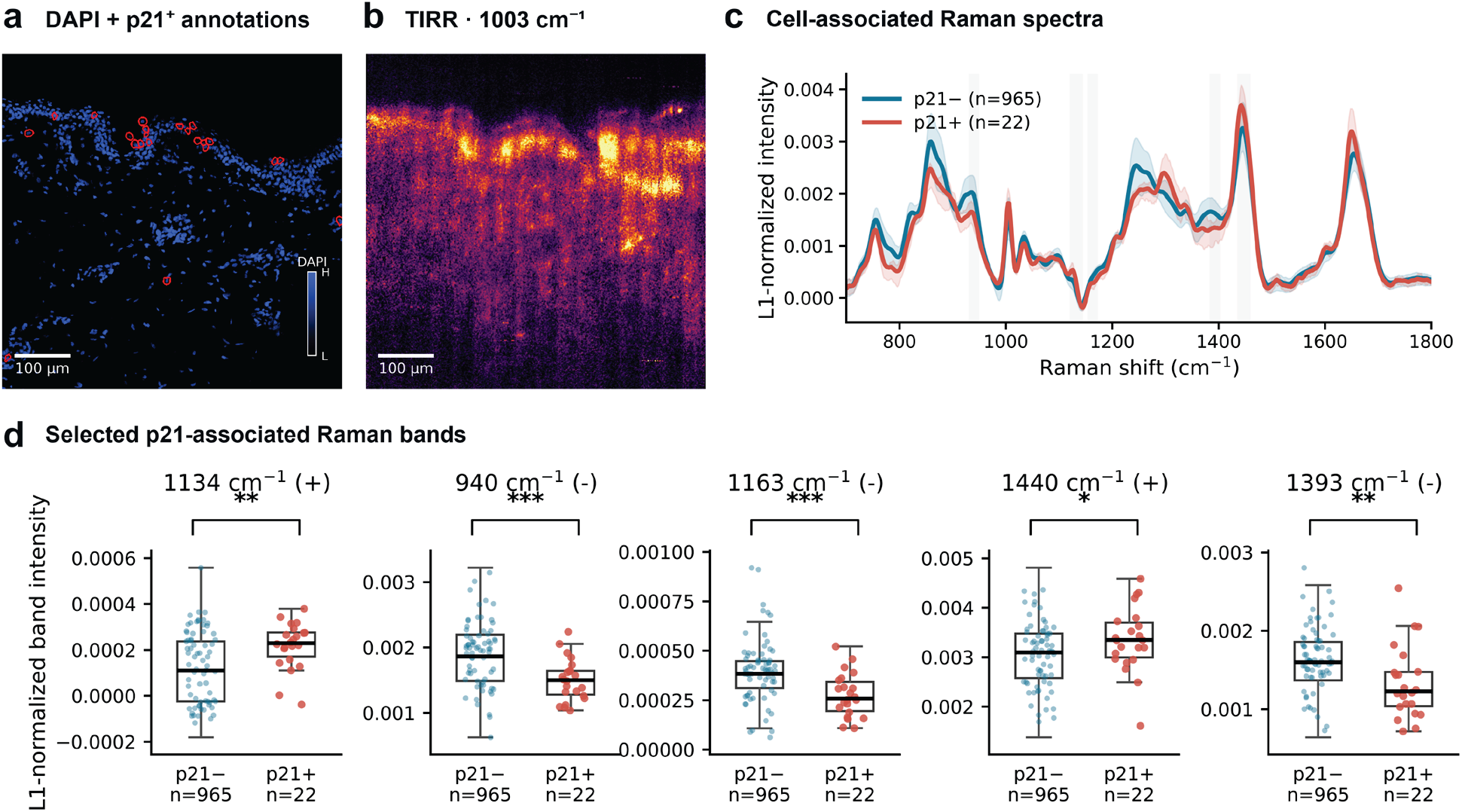
Integration of TIRR hyperspectral imaging with spatial molecular annotations of cellular senescence in human skin. (a) Registered DAPI fluorescence image (blue) of a human skin tissue section, with p21+ cells identified by STARmap-based in situ hybridization highlighted by red outlines. DAPI fluorescence intensity is displayed on a relative scale from low (black) to high (blue). (b) Corresponding TIRR Raman intensity map reconstructed near 1003 cm⁻¹ (phenylalanine), showing the spatial organization of the investigated tissue region. Scale bars, 100 µm. (c) Mean L1-normalized Raman spectra extracted from p21+ (red, n = 22) and p21− (blue, n = 965) cell-associated regions. Shaded areas surrounding the curves represent the interquartile range (25th–75th percentiles), while vertical gray bands indicate the spectral regions selected for quantitative comparison. (d) Distributions of normalized Raman band intensities near 1134, 940, 1163, 1440 and 1393 cm⁻¹ in p21+ and p21− cells. Boxes indicate the interquartile range, central lines denote the median, and whiskers extend to the most extreme observations within 1.5 times the interquartile range. Individual points represent cell-associated Raman measurements; p21− points were subsampled for visualization, while all eligible cells were included in the statistical analysis. Differences were evaluated using two-sided Mann–Whitney U tests, with p-values adjusted for five comparisons using the Holm procedure. Significance levels are indicated as *adjusted p < 0.05, **adjusted p < 0.01 and ***adjusted p < 0.001. The analysis represents one tissue section from a single human donor.

Comparison of the resulting spectra revealed distinct biochemical differences between p21+ and p21− populations across multiple vibrational regions (Fig. 6c). Guided by senescence-associated Raman signatures previously identified in mouse skin using the RamanOmics framework^35^, we examined five selected spectral bands (Fig. 6d). The p21+ population exhibited significantly higher normalized intensities near 1134 and 1440 cm⁻¹, associated with lipid-related vibrations, with rank-biserial correlation coefficients (RBCs) of +0.40 and +0.27, respectively. Conversely, reduced intensities were observed near 940 and 1163 cm⁻¹, associated with collagen-related spectral features (RBC = −0.46 and −0.49), and near 1393 cm⁻¹, associated with amino-acid vibrations (RBC = −0.41). All five differences remained statistically significant following Holm correction for multiple comparisons (adjusted p < 0.05), with detailed results reported in Supplementary Table 2. Notably, the direction of these changes closely parallels the RamanOmics signatures previously associated with senescence in aged mouse skin, despite the different tissue species and Raman sampling geometries. The concurrent enhancement of lipid-associated features and reduction of collagen- and amino-acid-associated signals suggests a redistribution of the relative biochemical contributions within p21-expressing cellular regions, potentially reflecting aspects of the lipid remodeling and extracellular-matrix alterations associated with cellular senescence. While these observations remain exploratory and are limited to a single human tissue section, their agreement with previously reported senescence-associated spectral trends suggests that biologically informative vibrational signatures can be retained under evanescent excitation. Beyond demonstrating molecular contrast in heterogeneous tissues, these results provide an initial proof of concept for coupling high-throughput TIRR imaging with spatial molecular profiling to investigate cellular phenotypes and their associated biochemical microenvironments.

## DISCUSSION

TIRR combines repeated evanescent excitation within a thin sapphire substrate with Hadamard-coded detection to decouple the illuminated area from the spatial detail recovered in the image. Unlike parallel illumination schemes that distribute incident power among separate spots, successive internal reflections extend excitation along the specimen interface. Although total internal reflection Raman spectroscopy is established for surface-sensitive measurements^16^, the present architecture couples this excitation mechanism to programmable hyperspectral detection. Despite an excitation width of approximately 50 µm, the reconstructed images exhibited effective 10–90% transition widths of approximately 3.5 µm. The coded aperture increased the spectral linewidth from approximately 7 cm⁻¹ with the conventional slit to approximately 10 cm⁻¹, a modest trade-off that remains compatible with recovery of characteristic tissue Raman bands. Together, these observations illustrate how extended illumination and computational spatial discrimination can be designed jointly to improve Raman imaging throughput.

The hyperspectral measurements of human skin and colon demonstrate the applicability of this principle to biological tissues with distinct architectures. TIRR recovered recognizable tissue organization and heterogeneous Raman contrast across submillimeter-scale fields of view, including enhanced signal near the presumed dermal–epidermal junction in skin and spatially distributed vibrational features in colon. Importantly, complete Raman fingerprints were retained at individual spatial locations rather than limiting acquisition to selected molecular bands. This capability is particularly relevant for investigating biological states characterized by complex spectral signatures.

The confocal benchmark demonstrates the practical throughput advantage of TIRR for the implementations tested. The observed time reduction was obtained at lower reported excitation power, although differences in effective spatial resolution, axial sampling and image processing limit the interpretation of the normalized ratio. The most direct conclusion is that TIRR can acquire spatially resolved tissue Raman spectra more rapidly under the stated experimental conditions.

Reduced-pattern reconstruction provides a second route to faster acquisition. The fixed 30-pattern low-sequency subset retained high agreement with the full-pattern reference in both the colon development dataset and the independent skin dataset. This result supports reuse of the selected masks for the two tissue architectures investigated, while the limitations of regularization are discussed below.

These two sources of throughput improvement are complementary. The experimentally derived 20.1-fold normalized acquisition-efficiency advantage was obtained with the complete 63-pattern sequence, whereas selecting 30 low-sequency patterns reduces the nominal integration budget by an additional factor of 63/30 = 2.1. Combining these factors suggests that a power- and SNR-normalized throughput enhancement approaching 40-fold could be attainable, corresponding to a projected factor of approximately 42 under the investigated conditions. Notably, this additional acceleration requires no modification of the optical hardware, relying instead on informed mask selection and computational reconstruction. Because the compressed reconstructions were evaluated retrospectively from complete acquisitions, the combined speedup remains a projection rather than a directly measured end-to-end improvement at matched spectral SNR. Its realization will depend on the noise characteristics of shortened acquisitions, instrumental overhead and the fidelity required for the intended application. Nevertheless, the independent tissue results demonstrate that a substantial reduction in measurement budget can preserve the principal spatial and vibrational information recovered by full-pattern imaging.

An important implication is that this increased throughput could make Raman-based molecular phenotyping more practical at tissue scales. By retaining multidimensional vibrational fingerprints, TIRR may support computational inference of biological states that cannot be identified from individual molecular bands. Previous work has demonstrated the possibility of relating Raman spectra to gene-expression profiles through approaches such as Raman2RNA^6^. Combining such methods with faster spatially resolved Raman acquisition could facilitate the investigation of molecular heterogeneity across larger tissue regions, linking label-free chemical imaging with emerging computational molecular profiling approaches.

The integration of TIRR with spatial molecular profiling provides an initial demonstration of its potential for investigating biologically defined cellular states. By associating Raman spectra with p21 expression in human skin, we identified spectral differences consistent with senescence-associated signatures previously reported using confocal Raman microscopy in mouse skin^35^. This agreement across distinct Raman imaging geometries and biological specimens suggests that at least some senescence-associated biochemical features may be preserved under surface-sensitive evanescent excitation. Although the present analysis is limited to a single donor and does not establish independent Raman-based identification of senescent cells, it highlights the potential of TIRR to investigate molecular heterogeneity associated with cellular phenotypes. With further biological validation and improved spatial registration, the combination of accelerated hyperspectral imaging and molecular annotations could provide a valuable framework for studying the biochemical microenvironment of senescent cells and other biologically relevant cell populations across larger tissue regions.

Despite the reported advantages, some limitations remain relevant to further development of the platform. The evanescent excitation geometry imposes a requirement for close specimen–substrate contact, making sample preparation and interfacial uniformity important determinants of imaging performance. The surface-sensitive nature of the measurement also restricts the depth from which Raman information is collected, which may be advantageous for interrogating well-defined interfaces but limits direct generalization to thicker or volumetric specimens. In addition, the spatial resolution of the present implementation is influenced by the complete imaging and encoding pathway, including relay optics, DMD sampling and numerical reconstruction, rather than by the collection objective alone. The spectral response is similarly affected by the coded-aperture geometry, requiring correction of position-dependent spectral displacements and careful characterization of the resulting spectral resolution. Finally, although compressive reconstruction preserves much of the information present in the full-pattern datasets, regularization can suppress weak or highly localized spatial and spectral features. High SAM and SSIM agreement therefore provides evidence of reconstruction fidelity but does not guarantee preservation of every chemically or biologically informative signal. These considerations underscore the importance of evaluating compressed acquisition not only using global reconstruction metrics but also through application-specific biological endpoints.

Future improvements could address several of these constraints without altering the fundamental principle of TIRR. Optimizing the substrate geometry and optical coupling may improve the uniformity of evanescent excitation, while refinements to the detection optics and DMD sampling could increase spatial resolving performance. Faster detector readout and reduced acquisition overhead would further increase the benefit of collecting fewer patterns. On the computational side, more advanced spatial–spectral priors, calibration-aware reconstruction and adaptive pattern selection may permit additional reductions in the measurement budget, particularly when the spectral or morphological characteristics of the specimen are known in advance. The programmable nature of the detection architecture makes these developments especially attractive, as acquisition strategies could be adjusted to prioritize spectral fidelity, spatial resolution or throughput depending on the biological question. Ultimately, TIRR illustrates a broader strategy for spontaneous Raman microscopy in which illumination geometry, spatial encoding and computational reconstruction are designed jointly rather than optimized independently. By combining extended evanescent excitation with hyperspectral detection and informed measurement selection, this framework offers a path toward substantially faster, label-free molecular imaging of biological tissues while retaining the rich spectral information needed for increasingly sophisticated computational analyses.

## MATERIALS AND METHODS

### Optical setup

The TIRR microscope was built around a 785 nm continuous-wave diode laser (Cobolt 08-01, 500 mW nominal output power, Hübner Photonics). The excitation beam was focused using an aspheric lens (f = 11 mm) and coupled into a polished sapphire substrate (10 × 10 mm², approximately 30 µm thick, Valley Design Corp.) through a 2 mm uncoated right-angle prism made of high-refractive-index N-SF11 glass (stock #45-524, Edmund Optics). At the excitation wavelength, the refractive index of N-SF11 (n ≈ 1.77) closely matches that of sapphire (n ≈ 1.76). A high-refractive-index immersion oil (n ≈ 1.64) was introduced between the prism and sapphire surfaces to minimize refractive-index discontinuities and Fresnel reflection losses, thereby facilitating efficient optical coupling into the substrate. Following coupling, the laser beam propagated laterally through successive total internal reflections between the sapphire interfaces, producing evanescent excitation at the specimen–substrate boundary.

Raman-scattered light was collected through the sapphire substrate using a 40×/0.95 NA dry objective (UPLXAPO40X, Olympus), providing a nominal field of view of approximately 660 µm. Elastically scattered excitation light was rejected using a dichroic mirror (LPD02-785RU, Semrock) and a long-pass edge filter (LP02-785RE, Semrock). The collected Raman image was subsequently relayed onto a digital micromirror device (DLP7000, Texas Instruments) positioned at an intermediate conjugate image plane. The DMD comprised an array of 1024 × 768 independently addressable micromirrors with a pitch of 13.68 µm and a nominal tilt angle of ±12°. The device was configured as a programmable reflective aperture, with micromirrors in the on state directing selected spatial components of the Raman signal toward the spectrograph, while those in the off state redirected the corresponding light away from the detection path. Binary spatial encoding patterns were sequentially displayed on the DMD to enable multiplexed spectral acquisition, as described in the following section.

The spatially encoded Raman signal was relayed into an imaging spectrograph (LS-785, Princeton Instruments; f/2, 85 mm focal length), where it was spectrally dispersed onto a back-illuminated CCD camera (PIXIS:400, 1340 × 400 pixels, Princeton Instruments). The acquisition configuration covered an approximate Raman-shift interval of 600–2200 cm⁻¹, corresponding to a spectral sampling interval of approximately 1.2 cm⁻¹ per detector pixel. The experimentally measured spectral resolving performance of the system was characterized separately using narrow emission lines from a Ne–Ar calibration source.

### Tissue sample preparation

All human tissue specimens were obtained and used in accordance with applicable institutional ethical guidelines and regulations. Human skin and colon biopsy specimens were stored at −80 °C until processing. Frozen tissue samples were embedded in optimal cutting temperature (OCT) compound and sectioned using a cryostat to obtain 14-µm-thick tissue sections. The sections were transferred onto polished sapphire substrates approximately 30 µm thick, which had been pretreated with poly-L-lysine (PLL) to promote tissue adhesion. This surface treatment was particularly important for the TIRR illumination geometry, as Raman excitation is generated by the evanescent field at the sapphire–specimen interface. Efficient excitation therefore depends on maintaining close contact between the tissue and the substrate, minimizing interfacial gaps that would otherwise attenuate the evanescent field before reaching the specimen.

Following transfer onto the sapphire substrates, the tissue sections were fixed with 4% paraformaldehyde (PFA) for 10 min. The specimens were subsequently rinsed with distilled water to remove residual fixative and minimize the deposition of buffer salts during drying. After rinsing, the samples were allowed to dry and all TIRR measurements were performed under dry conditions. This preparation procedure was used for both skin and colon specimens to maintain tissue adhesion and provide consistent conditions for evanescent-field Raman excitation.

### Binary Hadamard coded-aperture encoding and reconstruction

Spatial encoding of the Raman signal was implemented using a digital micromirror device (DMD) programmed with binary patterns derived from a Hadamard matrix^20,36^. In the configuration used here, a Hadamard matrix of order 64, **H** ∈ {−1, +1}^64×64^, was generated and converted into a binary encoding matrix by replacing +1 with 0 and −1 with 1. The first row and column were subsequently removed, yielding a 63 × 63 binary matrix **S**:

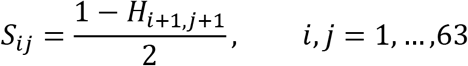

Unlike conventional Hadamard matrices, whose entries are ±1, the resulting S-matrix contains only 0 and 1, corresponding to the closed and open states of the DMD aperture. Each row of **S** defines an acquisition pattern, with 32 of the 63 encoded spatial elements transmitting light toward the spectrograph^17^. The complete set of 63 patterns provides a square, invertible measurement matrix, although the binary transformation does not preserve the orthogonality of the original Hadamard basis. The consequences of this distinction for multiplexing performance and noise propagation are discussed separately in Supplementary Note 4.

To implement the spatial encoding experimentally, the binary values of the encoding matrix were mapped onto the DMD using logical pixels composed of 2 × 2 micromirrors. The Hadamard patterns were replicated along the appropriate diagonal coordinates of the DMD to encode the spatial distribution of Raman emission while preserving its correspondence with the detector coordinates.

To mitigate time-dependent signal variations associated with photobleaching of endogenous fluorophores in biological specimens, a temporally symmetric acquisition sequence was implemented. Rather than recording the 63 Hadamard patterns only once in sequential order, each complete pattern set was acquired in both forward and reverse order (1, 2, …, 63, 63, …, 2, 1), resulting in 126 CCD frames per acquisition tile. For the tissue datasets in Fig. 3, each frame was integrated for 40 s; corresponding forward and reverse measurements were combined to yield 80 s per effective Hadamard pattern.

This acquisition strategy was designed to reduce systematic biases arising from the gradual decay of fluorescence background during prolonged laser exposure. Because the two measurements associated with each Hadamard pattern are acquired symmetrically around the midpoint of the acquisition sequence, their combination compensates, to first order, for approximately linear temporal signal drift. This reduces the correlation between acquisition order and photobleaching-induced intensity variations, which could otherwise introduce pattern-dependent offsets in the coded measurements and propagate into spatial artifacts during reconstruction. Before reconstruction, isolated positive spectral spikes were suppressed using a nine-point median filter, replacing spectral samples exceeding the local median by more than 90 detector counts. The resulting measurements were used to reconstruct the spatially resolved Raman signals.

For a given detector line, the forward measurement model can be written as

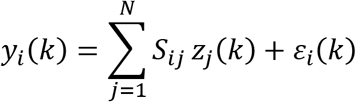

where *y_i_*(*k*) is the intensity recorded at detector spectral channel (k) for the (i)-th pattern, *S_iii_* is the binary encoding coefficient, and *z_i_* (*k*) is the spectrum associated with spatial element j, expressed in detector coordinates. The term *ε_i_*(*k*) accounts for measurement noise. Importantly, the spectra *z_i_* may be spatially shifted along the detector spectral axis; this effect is addressed independently by the spectral alignment procedure described below.

In matrix form, the measurement model reduces to

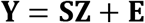

For full-pattern acquisition (N=63), the spatially resolved Raman spectra were recovered by solving the linear least-squares problem:

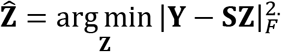

Because the binary encoding matrix S is square and full rank, the solution is unique and can be obtained without additional regularization. Minimization of the least-squares objective yields the normal equations:

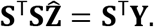

Rather than explicitly computing S^−1^, the system was solved numerically using a sparse direct solver. Independent reconstruction operators were applied to each detector row, neglecting contributions from adjacent rows.

### Hyperspectral cube assembly and preprocessing

Following the linear reconstruction of the coded-aperture measurements, the recovered Raman spectra were assigned to the corresponding spatial coordinates according to the diagonal mapping imposed by the DMD encoding geometry. This procedure generated an intermediate hyperspectral representation in which each reconstructed spatial element retained its complete Raman spectrum in detector coordinates. To recover the physical orientation of the specimen, the reconstructed array was geometrically rotated by approximately 46°, spatially cropped to exclude regions outside the valid reconstruction domain, and resampled using bilinear interpolation onto a rectangular image grid. In this configuration, each reconstructed tile comprised approximately 552 × 43 spatial pixels, with the Raman spectrum retained along the third dimension. Geometrical interpolation was performed independently for each spectral channel.

Spectral preprocessing was subsequently applied to the reconstructed hyperspectral tiles. Individual spectra were smoothed using a Savitzky–Golay filter with an 11-point window and a third-order polynomial. Fluorescence and slowly varying background contributions were estimated and removed using asymmetric least-squares baseline correction, with a smoothing parameter of λ = 10⁵, an asymmetry parameter of p = 0.01, and 10 iterations. Baseline estimation was performed independently for each reconstructed spatial spectrum, preserving local spectral variations while reducing non-Raman background contributions.

To generate extended hyperspectral images, the reconstructed tiles were assembled according to their acquisition order and the corresponding motorized-stage positions. Adjacent tiles were acquired with a nominal spatial overlap of 10 µm. Overlapping regions were combined using a linear cross-fade over eight reconstructed pixels, with the contribution from the preceding tile decreasing linearly as that from the subsequent tile increased. The same blending weights were applied independently at every spectral channel, producing a continuous hyperspectral mosaic without introducing wavelength-dependent stitching operations. The resulting three-dimensional dataset contained a complete reconstructed Raman spectrum at each spatial location.

### Pixel-wise spectral shift correction

In coded-aperture spectral imaging, light originating from different spatial positions enters the dispersive spectrograph at different entrance coordinates, resulting in position-dependent spectral displacements on the detector^20,23^. This spatial–spectral coupling can be described by a transformation acting independently on the Raman spectrum associated with each spatial element.

Let *x_i_* denote the Raman spectrum originating from spatial element *j*, and *S_ij_* ∈ 0,1 the corresponding binary encoding coefficient in the *i*-th DMD pattern. In the absence of spatially dependent spectral displacement, each measurement would be the sum of the spectra transmitted by the active DMD elements. To account for the spectrograph response, we introduce a position-dependent transformation *f_i_*, which maps the spectrum of each spatial element onto the corresponding detector coordinates. Assuming linear intensity response and a transformation independent of the applied DMD pattern, the recorded signal can be expressed as

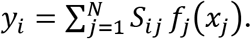

Defining the transformed spectrum as

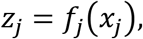

the measurement model reduces to the conventional coded-aperture formulation

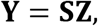

where **Y** contains the acquired measurements, **S** is the binary encoding matrix, and **Z** contains the spatially displaced spectra. For a complete set of linearly independent patterns, the individual spectra can therefore be recovered in detector coordinates through inversion of the encoding operation. In the ideal noise-free case, Z = S^−1^**Y**, whereas the experimental reconstruction employs the least-squares procedure described in the preceding section. Once the spatial contributions have been separated, the spectral transformation can be corrected independently for each reconstructed spatial element:

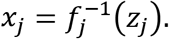

The dominant effect of the spatial–spectral coupling is a displacement of the Raman spectrum along the dispersion axis. Accordingly, the transformation *f_ii_* can be represented, to first order, as a position-dependent spectral translation:

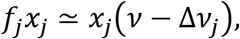

where *ν* denotes the Raman shift and Δ*ν_i_* is the displacement associated with spatial element *j*. This representation assumes that the relative displacement between spectral features remains approximately constant across the spectral range of interest.

To validate the rigid-shift approximation experimentally, we characterized the position-dependent spectral displacement across 31 DMD positions spanning the encoding region, using the central position as the reference. Spectral shifts were independently estimated through normalized cross-correlation in three spectral regions near 750, 1001 and 1602 cm⁻¹. The measured displacements exhibited an approximately linear dependence on DMD position and remained closely consistent across the three spectral regions (Supplementary Fig. S2). These measurements experimentally support the use of a rigid spectral translation to describe the dominant spatial–spectral coupling in the TIRR detection system.

In the experimental reconstruction, the spectral displacement of each spatial element was determined using the characteristic Raman band of the sapphire substrate at 650 cm⁻¹ as an internal reference^37^. Following coded-aperture reconstruction and image assembly, the sapphire peak was identified independently in each reconstructed spectrum within the 600–700 cm⁻¹ interval and aligned to a common reference position, conventionally set to 650 cm⁻¹. This internally referenced procedure allows spectral displacements to be corrected directly from the acquired Raman data, without relying on a fixed calibration of the DMD–spectrograph geometry, thereby simplifying spectral registration and reducing its dependence on the precise optical alignment.

### Comparison with confocal Raman microscopy

The spectral acquisition efficiency of TIRR was evaluated against conventional confocal Raman microscopy using hyperspectral measurements acquired from the same human colon tissue specimen over a corresponding tissue region. Both measurements employed near-infrared excitation at 785 nm. The TIRR dataset was reconstructed using the complete set of 63 Hadamard patterns and the reconstruction and preprocessing procedures described above, without compressive subsampling. The confocal dataset was acquired using point-scanning Raman microscopy, with the optical configuration and acquisition parameters detailed in Supplementary Note 3. The corresponding hyperspectral images were oriented and spatially cropped to identify the overlapping tissue region. The spatial sampling interval used for the comparison was approximately 1.2 µm in both datasets. Acquisition times and excitation powers were recorded for each measurement, yielding 3.1 h at 118 mW for TIRR and 27.9 h at 380 mW for confocal Raman microscopy.

Spectral SNR was calculated from the processed hyperspectral datasets using the phenylalanine Raman band near 1003 cm⁻¹. The corresponding signal intensity at each spatial pixel was defined as the mean spectral intensity within an approximately 10 cm⁻¹ window centered on the phenylalanine feature. Noise was estimated independently at each pixel as the standard deviation of the spectral intensity within a peak-free interval spanning approximately 1847–1870 cm⁻¹. The same spectral intervals, expressed in Raman-shift units, were used for both modalities, with the corresponding detector-channel ranges identified from their individual wavelength calibrations. The analysis was performed on the baseline-corrected and spectrally smoothed datasets, with Savitzky–Golay filtering applied using comparable spectral window widths to account for differences in spectral sampling^38^. The same processed spectra were used for both signal and noise estimation.

For a spatial location (p), the operational spectral SNR was defined as

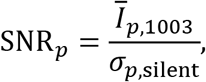

where I_p,1003_ is the mean intensity of the phenylalanine band and σ_p,silent_ is the spectral standard deviation within the peak-free reference interval. To exclude substrate regions and pixels with insufficient Raman signal, tissue masks were generated independently for the two modalities by thresholding their respective phenylalanine intensity maps and only pixels with finite SNR values were retained. The median of the resulting pixel-wise SNR distribution was used as the representative spectral SNR for each modality. The distributions were visualized using box plots showing the interquartile range, median and 5th–95th percentile whiskers.

Acquisition efficiency was quantified using both the measured acquisition-time ratio and a power- and SNR-normalized speedup factor. The latter was defined as

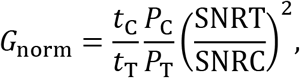

where the subscripts T and C denote TIRR and confocal microscopy, respectively, *t* is the acquisition time and *P* is the excitation power. Equivalently, this factor represents the ratio of the experimentally derived spectral acquisition efficiencies, defined as squared SNR per unit excitation power and acquisition time. Its interpretation as an equivalent imaging speedup at matched power and SNR assumes the conventional scaling SNR^2^ ∝ *Pt*. The comparison therefore characterizes acquisition efficiency under the investigated experimental and preprocessing conditions and does not impose equivalent spatial resolution or axial sampling depth between the two modalities.

### Compressive coded-aperture reconstruction

To investigate reconstruction from incomplete Hadamard measurements, subsets of *m* < 63 encoding patterns were selected from the full binary sensing matrix S. The corresponding reduced measurement model was expressed as

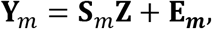

where S*_m_* ∈ 0,1*^m^*^×63^ contains the selected rows of the original encoding matrix, **Y***_m_* represents the acquired multiplexed measurements, and Z contains the unknown spatially resolved Raman spectra. Because the number of measurements is smaller than the number of encoded spatial elements, the reconstruction problem is underdetermined and requires additional constraints.

We addressed this problem through joint spatial–spectral total variation (TV) regularization, exploiting the spatial continuity of tissue structures and the correlation between neighboring spectral channels^39^. The hyperspectral reconstruction was formulated as the constrained optimization problem

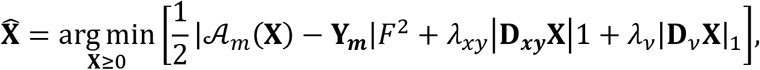

where A_m_ denotes the reduced Hadamard measurement operator applied to the hyperspectral image, D*_xy_* represents spatial finite differences, and D*_ν_* describes differences between adjacent spectral channels. Spatial regularization was implemented using horizontal and vertical nearest-neighbor connections derived from the physical pixel positions after diagonal DMD coordinate mapping, rather than from the ordering of the encoded elements alone. The resulting anisotropic TV penalty promotes spatial continuity while allowing intensity discontinuities associated with tissue structures, whereas the spectral TV term favors locally smooth Raman profiles without restricting the reconstruction to predefined vibrational bands.

The optimization problem was solved using a first-order primal–dual algorithm^40^, with non-negativity enforced at each iteration. Both regularization coefficients were fixed at *λ_xy_* = *λ_ν_* = 0.02 for normalized input measurements, with a maximum of 700 iterations and a relative convergence tolerance of 2 × 10^−5^. The same reconstruction parameters were used throughout the pattern-selection experiments and the independent tissue evaluation. Following reconstruction, the resulting hyperspectral cubes were processed using the geometrical and spectral correction procedures described above.

### Low-sequency pattern selection and reconstruction fidelity assessment

The influence of Hadamard pattern selection was initially investigated by reconstructing a colon tissue dataset using randomly selected subsets containing 10, 20, 30, 40, 50 and 60 patterns. For each measurement budget, ten subsets were sampled without replacement from the complete set of 63 masks, and the corresponding hyperspectral images were reconstructed using the same joint spatial–spectral TV regularization.

To characterize the spatial modulation introduced by each mask, we calculated its sequency, defined as the number of transitions between adjacent binary elements:

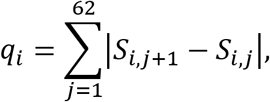

where *S_i_*_,*ii*_ is the binary value of the *j*-th encoded element in mask *i*. Low-sequency masks contain slowly varying spatial patterns, whereas high-sequency masks exhibit more rapid alternation between transmitting and non-transmitting elements. This distinction is analogous to spatial frequency ordering in the Walsh–Hadamard basis and has previously been investigated in compressive single-pixel imaging^41,42^.

For each randomly selected subset, we quantified its enrichment in low-sequency masks by comparing the mean sequency of the selected patterns with that of the complete encoding matrix. The resulting enrichment score was standardized using the expected sampling variability for random selection without replacement. To separate the influence of mask composition from that of measurement number, SAM and SSIM values were centered by subtracting the corresponding mean within each measurement budget, and Spearman correlations were calculated between these residuals and the low-sequency enrichment scores. Based on the observed associations, a deterministic subset was constructed by selecting the 30 patterns with the lowest sequency, resolving equal-sequency ties by their original matrix order. The resulting mask indices were fixed and subsequently applied without modification to an independent skin tissue dataset.

Reconstruction fidelity was evaluated relative to the corresponding complete 63-pattern hyperspectral reference using two complementary metrics. Spectral similarity was quantified using the spectral angle mapper (SAM), calculated between the reference and reconstructed spectra at corresponding spatial locations:

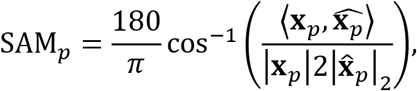

where **x***_p_* and **x**x*_p_* represent the reference and reconstructed Raman spectra at pixel *p*. The reported SAM was the median spectral angle over pixels exceeding a reference signal threshold corresponding to 5% of the 99th percentile of the reference spectral norms, excluding locations with vanishing reconstructed signal. Lower SAM values indicate stronger agreement in spectral shape.

Spatial fidelity was quantified using the SSIM applied to two-dimensional maps of the spectral Euclidean norm, *I*(*x*, *y*) = √∑*v X* (*x*, *y*, *v*)^2^. Reference and reconstructed maps were normalized using the same intensity scale derived from the reference image and evaluated within the bounding region of the reference signal mask. SSIM was calculated using Gaussian-weighted local statistics (*σ* = 1.5), with values approaching unity indicating stronger agreement in spatial structure. Unlike SAM, which primarily characterizes the relative distribution of spectral features, SSIM evaluates the preservation of spatial intensity patterns. Both metrics were calculated over the reconstructed hyperspectral datasets rather than exclusively from the Raman bands displayed in Fig. 5.

### STARmap-ISH for senescence analysis

After TIRR imaging, the same human skin section was processed by STARmap-based in situ hybridization (STARmap-ISH) to map *CDKN1A* transcripts, which encode p21, for the senescence analysis (Fig. 6)^34^. The protocol was adapted from Zhang et al.^35^. The section was permeabilized in prechilled methanol at −20 °C for 30 min, equilibrated to room temperature and treated with 10 mM Tris (pH 7.5) for 10 min. STARmap probes (Supplementary Table 3) were hybridized at 10 nM per probe for 48 h at 40 °C with gentle shaking. Probes were ligated with T4 DNA ligase and amplified by Phi29-mediated rolling-circle amplification. Amplicons were functionalized and embedded in a polyacrylamide gel, and the tissue–gel hybrid was cleared with proteinase K and dephosphorylated. For detection, gene-specific fluorescent oligonucleotides (Supplementary Table 3) were applied at 100 nM in 1× SSC in PBST for 90 min at room temperature, followed by three 5-min PBST washes. Nuclei were counterstained with DAPI, and fluorescence images were acquired on a Leica Stellaris 5 confocal microscope with a 405-nm diode laser, a white-light laser and a 20× air objective.

DAPI and gene-channel images were stitched from tiles acquired with 5% overlap, with an additional 5-pixel margin on each side. Fluorescent RNA spots were detected independently in each gene channel as local maxima in a 5 × 5-pixel neighborhood and retained when their local contrast exceeded a sample-specific threshold derived from the intensity distribution. Nuclei were segmented from DAPI images using the pretrained Cellpose-SAM model in Cellpose v4.0.4^43^. Nuclear masks were expanded by 3 pixels using label-preserving dilation to approximate whole-cell regions while preventing overlap. Each spot was assigned to a cell if the center of its bounding box fell within an expanded mask. Unassigned spots were excluded from downstream analysis.

### TIRR–DAPI spatial registration and cell-associated spectral extraction

Spatial registration between TIRR hyperspectral images and DAPI fluorescence microscopy was performed using a semi-automated landmark-guided approach. A DNA-associated Raman intensity map obtained by spectral fitting was used as the reference for alignment with the corresponding DAPI image. Two recognizable nuclear regions were manually selected in the TIRR image, together with their approximate matching regions in the DAPI mosaic. Following initial rescaling based on the physical pixel sizes of the two modalities, a joint optimization of rotation, translation and a small additional scaling correction was performed using a coarse-to-fine search. For each candidate transformation, normalized cross-correlation was calculated between local TIRR patches and their corresponding DAPI regions. The optimization maximized a combined similarity score defined as the mean of the two local correlation coefficients minus one-quarter of their absolute difference, favoring transformations that consistently aligned both regions. The resulting affine transformation was applied to the fluorescence-derived cell segmentation masks using nearest-neighbor interpolation, preserving individual cell labels. A mean Raman spectrum was subsequently extracted from the pixels associated with each registered cellular region.

### Statistical analysis of senescence-associated Raman signatures

Cell-associated Raman spectra were normalized to their L1 norm over the 600–1800 cm⁻¹ fingerprint region. Cells represented by fewer than three reconstructed TIRR pixels were excluded. Cells were classified as p21+ or p21− according to the corresponding STARmap molecular annotations. Raman band intensities were calculated as the Simpson-integrated mean intensity within five selected spectral intervals centered near 1134, 940, 1163, 1440 and 1393 cm⁻¹. Differences between p21+ and p21− populations were assessed using two-sided Mann–Whitney U tests^44^. P-values were adjusted across the five comparisons using the Holm procedure^45^. Effect sizes were expressed as rank-biserial correlation coefficients (RBC), with positive values indicating higher Raman intensities in p21+ regions. Statistical significance was defined as adjusted P < 0.05.

## ACKNOWLEDGMENTS

We thank colleagues at the Cutaneous Biology Research Center for providing resources and support. This work was supported by the National Institutes of Health (NIH) New Innovator Award (DP2TR004354; J.S.), the NIH Common Fund (UH3CA275687 and UG3CA275687; P.T.C.S., J.S., and J.W.K.), NIH grants P41EB015871 and P01AI129859 (the latter to J.S.), the National Cancer Center, Korea (NCC-24H1170), the MIT–MGB Seed Grant, the Massachusetts Life Science Center (J.S.), the Burroughs Wellcome Fund (J.S.), Additional Ventures (J.S.), and Massachusetts General Hospital (J.S.). L.B. was supported by the MIT/DFHCC Bridge Project Grant, the Margaret Q. Landenberger Research Foundation, the Cancer Research Institute, the Krantz Quantum Award, and the MGB Transformative Scholar Award.

## Author contributions

K.J.K.-K., P.T.C.S., J.W.K. and J.S. conceived the study. S.S. and K.J.K.-K. built and characterized the optical setup and developed the coded-aperture reconstruction software. S.S. optimized the optical setup, prepared the samples for Raman imaging and performed Raman measurements. S.S. and K.J.K.-K. developed the compressed sensing approach. K.Z. and H.H. performed STARmap profiling and segmentation for the senescence analysis. S.S. performed the data and statistical analysis, prepared the figures and wrote the original draft of the manuscript. P.T.C.S., K.J.K.-K., J.W.K., L.B. and J.S. secured funding for the study. P.T.C.S., J.W.K. and J.S. supervised the study. All authors reviewed and commented on the manuscript.

## Competing interests

A patent application covering aspects of the TIRR technology described in this work has been filed by MIT, with S.S., K.J.K.-K., P.T.C.S. and J.W.K. listed as inventors. J.S. serves as a scientific advisor to Johnson & Johnson and Aë Laboratories. The other authors declare no competing interests.

## Data and code availability

Representative Raman images and spectra, together with the quantitative results, are presented in the manuscript and Supplementary Information. The raw hyperspectral Raman data and the code used for coded-aperture reconstruction and compressed sensing are available from the corresponding authors upon reasonable request.

