## Supplementary information for "High-Throughput Coded Aperture Raman Imaging with Total Internal Reflection"

### Supplementary Note 1 | Experimental characterization of spectral resolution

The spectral resolving performance of the TIRR detection system was characterized by comparing narrow calibration-source emission features recorded under two optical configurations: conventional detection with the spectrograph slit closed and coded-aperture detection. Both spectra were represented using the same experimentally determined Raman-shift-equivalent calibration axis. The closed-slit acquisition contained 250 additional detector channels at the beginning of the recorded array, which were excluded to establish the corresponding spectral coordinate range.

Four relatively isolated spectral features were selected for the analysis, located near 640, 690, 800, and 1780 $cm^{-1}$ on the Raman-equivalent axis. Corresponding features were identified independently in the two acquisitions. A local linear baseline was defined for each peak using median intensities from two spectral intervals on either side of the feature. This approach accounts for local variations in spectral background without introducing assumptions about the functional form of the emission-line profile.

For a recorded spectrum $I(k)$ and its local baseline $B(k)$, the baseline-corrected peak height was defined as

$$H = I(k_{\max}) - B(k_{\max}),$$

where $k_{\max}$ denotes the detector channel containing the local maximum. The half-maximum level was then determined as (H/2). On each side of the peak, the detector sample nearest to this level was identified from the baseline-corrected intensity. The spectral linewidth was calculated directly as

$$\mathrm{FWHM} = |\nu(k_R) - \nu(k_L)|,$$

where $k_L$ and $k_R$ are the selected detector channels, and $\nu(k)$ is the calibrated spectral coordinate. No spectral smoothing or Gaussian/Lorentzian profile fitting was applied. The same analysis procedure was used for both optical configurations.

The four selected lines yielded FWHM values of 8.23, 6.06, 5.82, and 8.01 $cm^{-1}$ in the closed-slit configuration, compared with 10.31, 10.12, 11.67, and 8.04 $cm^{-1}$ under coded-aperture detection. The resulting mean linewidths were 7.03 ± 1.26 and 10.04 ± 1.50 $cm^{-1}$, respectively, where the reported standard deviations describe variations across the four investigated spectral features rather than independent experimental repetitions. The measured linewidths provide an empirical characterization of the spectral response under each detection configuration.

For visual comparison of the complete spectra in Supplementary Fig. S3a, a single rigid spectral offset of +7.20 $cm^{-1}$ was applied to the displayed coded-aperture coordinate axis.

This value corresponds to the mean difference between the four matched peak positions. The offset was used exclusively for visualization; all linewidth measurements were performed on the original detector samples before spectral registration. In the individual peak comparisons, spectra were centered on their respective maxima to facilitate direct assessment of the measured widths.

Because the linewidths were extracted from a finite number of detector samples, their numerical precision is limited by the local spectral sampling interval. Nevertheless, the analysis provides a direct comparison of the response obtained with conventional slit-based detection and coded-aperture acquisition, without introducing model-dependent assumptions about peak shape. The observed linewidths support an effective coded-aperture spectral resolution on the order of 10 $cm^{-1}$ across the investigated spectral positions.

**Supplementary Note 2 | Influence of sampling budget and Hadamard mask composition on reconstruction fidelity**

To characterize the performance of compressed TIRR reconstruction, we performed a systematic undersampling analysis using the complete 63-pattern acquisition of the colon tissue specimen. For each measurement budget of 10, 20, 30, 40, 50 and 60 patterns, ten subsets were randomly sampled without replacement from the same binary Hadamard encoding matrix. Each subset was reconstructed independently using joint spatial–spectral total variation (TV) regularization, with identical regularization coefficients and convergence settings. A reconstruction using all 63 patterns was also performed with the same TV algorithm to characterize its limiting performance. All reconstructed hyperspectral datasets underwent the same geometrical and spectral post-processing procedures before comparison with the conventional full-pattern reference.

The dependence of reconstruction fidelity on measurement number is summarized in Supplementary Fig. S6 and Supplementary Table 1. Spectral fidelity was quantified using the median pixel-wise spectral angle mapper (SAM), whereas spatial fidelity was evaluated using structural similarity (SSIM) between maps of the spectral Euclidean norm. Each randomly selected subset yielded one SAM and one SSIM value. The curves in Supplementary Fig. S6 represent the mean of these values across the ten trials at each sampling level, with error bars indicating their standard deviation. The overall trend showed progressively improved reconstruction fidelity as additional patterns were included, although substantial differences remained between subsets containing the same number of measurements. This variability was particularly evident at intermediate sampling levels, indicating that reconstruction quality could not be predicted from the number of acquired patterns alone.

We therefore examined whether the composition of the selected subsets could explain part of this variability. Each Hadamard pattern was assigned a sequency value corresponding to the number of transitions between consecutive binary elements in its encoding vector. Sequency was calculated along the one-dimensional ordering of the Hadamard matrix, rather than interpreted as a direct measurement of two-dimensional spatial Fourier frequency after DMD mapping. For each subset, we defined a low-sequency enrichment score from the difference between its mean sequency and the population mean, standardized by the finite-population standard error associated with sampling without replacement. Positive enrichment scores therefore indicate a greater representation of slowly varying patterns than expected under random selection.

As a complementary descriptor, patterns were divided into low-sequency (1–20 transitions), intermediate-sequency (21–41 transitions) and high-sequency (42–62 transitions) categories. The LOW/HIGH balance was defined as the fraction of selected low-sequency patterns minus the fraction of high-sequency patterns. This balance was also standardized relative to its expected value and sampling variability in the complete mask population. The two descriptors capture related aspects of mask composition: the former uses the complete sequency distribution, whereas the latter emphasizes the relative representation of the extreme sequency categories.

To compare subsets acquired with different measurement budgets, SAM and SSIM residuals were calculated by subtracting the corresponding mean metric at each mask count. Spearman rank correlations were then evaluated between these residuals and the standardized mask-composition descriptors (Supplementary Fig. S7). Low-sequency enrichment was associated with reduced SAM residuals ($\rho = -0.30$) and increased SSIM residuals ($\rho = +0.29$). Similar trends were observed for the LOW/HIGH balance ($\rho = -0.28$ and +0.27, respectively). These pooled correlations summarize the overall association after accounting for the mean dependence of reconstruction quality on measurement number.

Importantly, the strength of these associations varied with the sampling budget (Supplementary Fig. S8). At 30 and 40 patterns, low-sequency enrichment was consistently associated with lower spectral error and greater spatial similarity, with correlation magnitudes approaching 0.5–0.6. The relationships were less consistent at 20 patterns and weakened near the complete measurement budget, where reconstruction performance became less sensitive to which patterns were included. These observations motivated the deterministic selection of the 30 lowest-sequency patterns, corresponding to sequency values from 1 to 30 in the encoding matrix. This subset was constructed directly from the matrix structure, independently of the reconstruction metrics obtained for individual random trials.

The selected 30-pattern subset achieved a SAM of 7.37° and an SSIM of 0.8633 on the colon development dataset. For comparison, the best SAM among the ten randomly sampled 30-pattern subsets was 9.01°, while the highest SSIM was 0.7600; these two values were obtained from different random subsets. The fixed low-sequency selection was subsequently applied without modification to an independent skin tissue dataset, yielding a SAM of 7.24° and an SSIM of 0.8846, as shown in Fig. 5.

The 63-pattern TV-regularized reconstruction should be distinguished from the conventional complete-pattern reference used to calculate the evaluation metrics. Although both reconstructions use the same complete measurement dataset, they employ different inversion procedures. Consequently, their comparison gives nonzero SAM and SSIM below unity, whereas comparison of the reference against itself gives SAM = 0° and SSIM = 1. The mask-composition correlations should be interpreted as exploratory associations derived from repeated subsets of a single development dataset, rather than as independent biological replicates or evidence of a universal optimal ordering. Together with the independent skin reconstruction, they provide experimental support for a transferable, structurally informed measurement-selection strategy.

**Supplementary Note 3 | Confocal Raman microscopy setup and acquisition parameters**

Confocal spontaneous Raman measurements were performed using a custom-built near-infrared Raman microscope operating at an excitation wavelength of 785 nm. The excitation source was a continuous-wave diode laser (Cobolt 08-DPL), spectrally filtered using a laser line filter (LL01-785-12.5, Semrock) and directed through dual-axis galvanometric scanning mirrors (CT 6210, Cambridge Technology). Large-area sample positioning was controlled using an XY motorized stage (MS-2000, ASI Imaging). A 40×/0.95 NA objective (UPLSAPO40X2, Olympus) was used for both excitation focusing and collection of the backscattered Raman signal. A brightfield imaging path enabled visualization of the specimen and identification of the tissue region for subsequent Raman acquisition.

The collected Raman light was separated from the excitation beam using dichroic optics and coupled into a multimode optical fiber, providing spatial filtering for confocal detection. The signal was subsequently dispersed by an imaging spectrograph (HoloSpec f/1.8i, Kaiser Optical Systems) and detected using a thermoelectrically cooled, back-illuminated, deep-depletion CCD camera (PIXIS:100BR_eXcelon, Princeton Instruments). Instrument control, beam scanning and hyperspectral acquisition were performed using custom MATLAB software.

For the throughput comparison with TIRR presented in Fig. 4, a hyperspectral confocal Raman image of 552 × 120 pixels was acquired from the same human colon tissue specimen and corresponding spatial region. The spatial sampling interval was set to 1.2 μm in both dimensions, matching the nominal sampling interval of the TIRR reconstruction and corresponding to an imaged area of approximately 662 × 144 $\mu m^2$. The confocal measurement was performed at an excitation power on the sample plane of 380 mW, with a nominal spectral integration time of 1.2 s per pixel. The complete hyperspectral acquisition required 27.9 h, including the instrumental overhead associated with sequential point-scanning measurements. The resulting dataset was compared with the corresponding TIRR acquisition, which required 3.1 h at 118 mW, as described in the main Methods section.

**Supplementary Note 4 | Shot-noise considerations for Hadamard multiplexing**

Hadamard multiplexing can provide a signal-to-noise ratio (SNR) advantage when detector read noise dominates the measurement. However, under shot-noise-limited conditions, the increased photon flux associated with multiplexed detection also increases the measurement variance. To clarify this distinction for the present TIRR architecture, we consider the noise propagation associated with reconstruction using the binary Hadamard encoding matrix.

For a single spectral channel, the multiplexed measurement model is

$$\mathbf{y} = \mathbf{S}\mathbf{x} + \mathbf{n},$$

where $\mathbf{x}$ contains the Raman intensities of the encoded spatial elements, $\mathbf{S}$ is the binary encoding matrix and $\mathbf{n}$ represents measurement noise. The experimental matrix was constructed by converting a Hadamard matrix of order 64 into binary values and removing its first row and column, yielding $\mathbf{S} \in 0,1^{63\times 63}$. Each encoding pattern contains 32 transmitting elements, and the matrix satisfies

$$\mathbf{S}^T\mathbf{S} = 16(\mathbf{I} + \mathbf{J}),$$

Where $\mathbf{I}$ is the identity matrix and $\mathbf{J}$ is the all-ones matrix.

For analytical simplicity, assume that all 63 spatial elements contribute the same mean signal $x$, expressed in detected photoelectrons per exposure, and that independent Poisson shot noise dominates. Each multiplexed measurement then has mean intensity and variance

$$\mathbb{E}[y_i] = 32x, \qquad \mathrm{Var}(y_i) = 32x.$$

The reconstructed spatial intensities are obtained using the least-squares estimator

$$\hat{\mathbf{x}} = (\mathbf{S}^T\mathbf{S})^{-1}\mathbf{S}^T\mathbf{y}.$$

Propagation of the measurement covariance through this linear reconstruction gives

$$\mathrm{Cov}(\hat{\mathbf{x}}) = 32x(\mathbf{S}^T\mathbf{S})^{-1} = 2x\left(\mathbf{I} - \frac{\mathbf{J}}{64}\right).$$

Consequently, the reconstructed variance of each spatial element is

$$\mathrm{Var}(\hat{x}_j) = \frac{63}{32}x.$$

For comparison, a direct measurement of one spatial element with the same exposure duration has shot-noise variance $x$ and SNR equal to $\sqrt{x}$. The ratio between the reconstructed multiplexed SNR and the direct-measurement SNR is therefore

$$\sqrt{\frac{32}{63}} \approx 0.71.$$

Thus, for uniformly emitting spatial elements, the binary S-matrix does not provide an intrinsic per-element SNR advantage in the shot-noise-limited regime. For reference, an ideal signed Hadamard matrix with orthogonal ±1 elements yields an SNR ratio of unity under the corresponding idealized shot-noise model. Such signed measurements, however, require a differential or complementary optical implementation and are not equivalent to the single binary-mask exposures used here.

For spatially heterogeneous Raman signals, the reconstructed noise covariance depends on the distribution of intensities across the encoded elements, and the uniform-signal expression is no longer generally applicable. Additional detector noise, fluorescence background and other experimental contributions may further modify the multiplexing behavior.

Importantly, the throughput improvement demonstrated by TIRR does not rely on a theoretical Hadamard SNR advantage. Instead, it originates from extended evanescent excitation and the parallel collection of spatially encoded Raman signals. The binary Hadamard matrix enables their computational separation, while the subsequent compressed reconstruction reduces the number of required measurements through spatial–spectral regularization and informed pattern selection.

**Supplementary Figure 1**

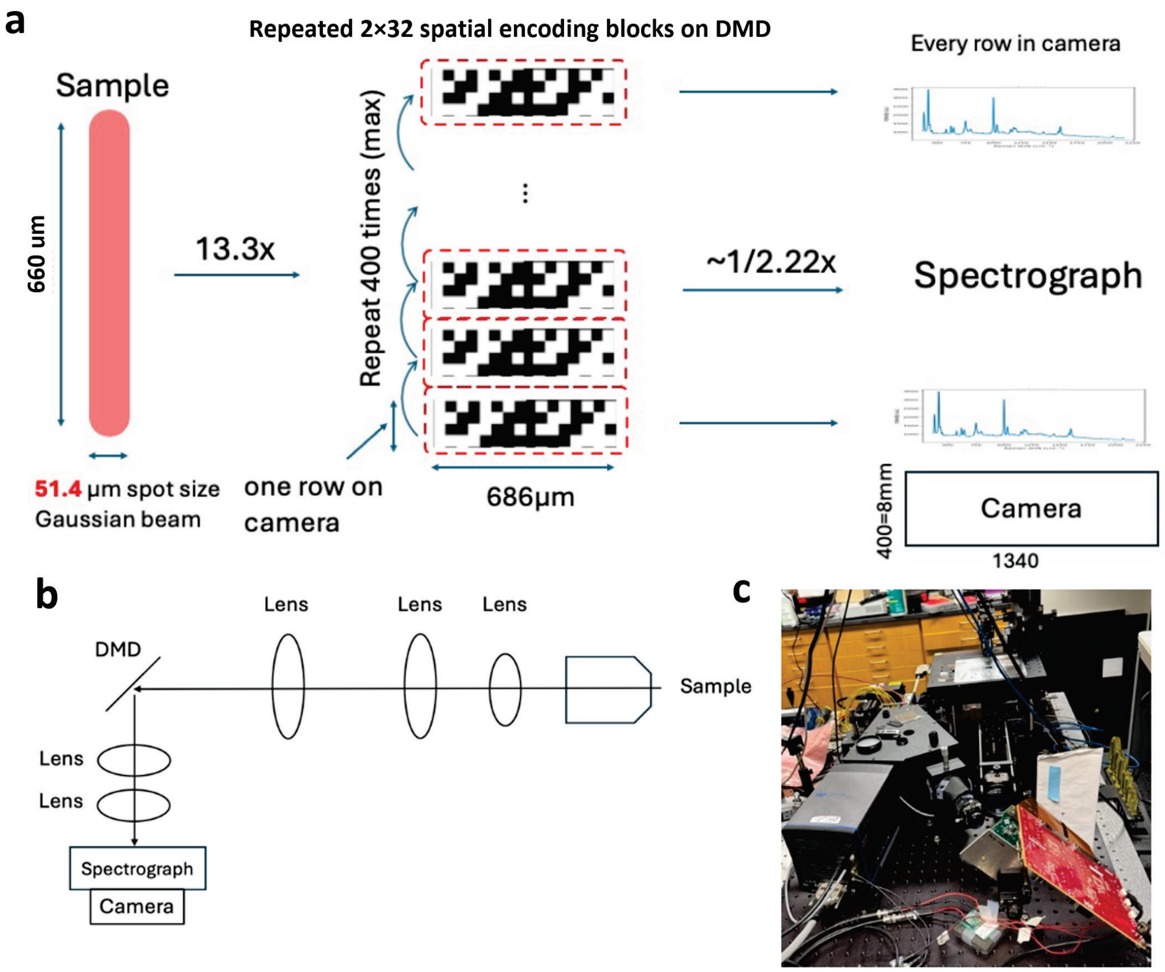


**Supplementary Figure 1 | Coded-aperture hyperspectral detection principle and experimental implementation in TIRR microscopy. (**a) Schematic representation of spatial encoding in the TIRR detection arm. Raman emission from the extended excitation region is projected onto a digital micromirror device (DMD), where repeated binary patterns select spatial contributions directed toward the spectrograph. The encoding patterns are arranged along diagonal DMD coordinates to maintain the correspondence between the encoded spatial positions and detector rows. In the experimental configuration, each logical encoding element comprises 2 × 2 micromirrors, and a complete measurement sequence consists of 63 binary Hadamard patterns, each containing 32 transmitting elements. (b) Simplified optical layout of the detection arm, showing image relay from the specimen onto the reflective DMD, followed by projection of the encoded Raman signal into the dispersive spectrograph and CCD camera. (c) Photograph of the laboratory optical arrangement. In contrast to conventional single-position Raman detection, the coded-aperture configuration collects multiplexed Raman signals from several spatial elements within each exposure. The individual spatial spectra are subsequently recovered by inversion of the corresponding encoding matrix, with position-dependent spectral displacements corrected after reconstruction.

**Supplementary Figure 2**

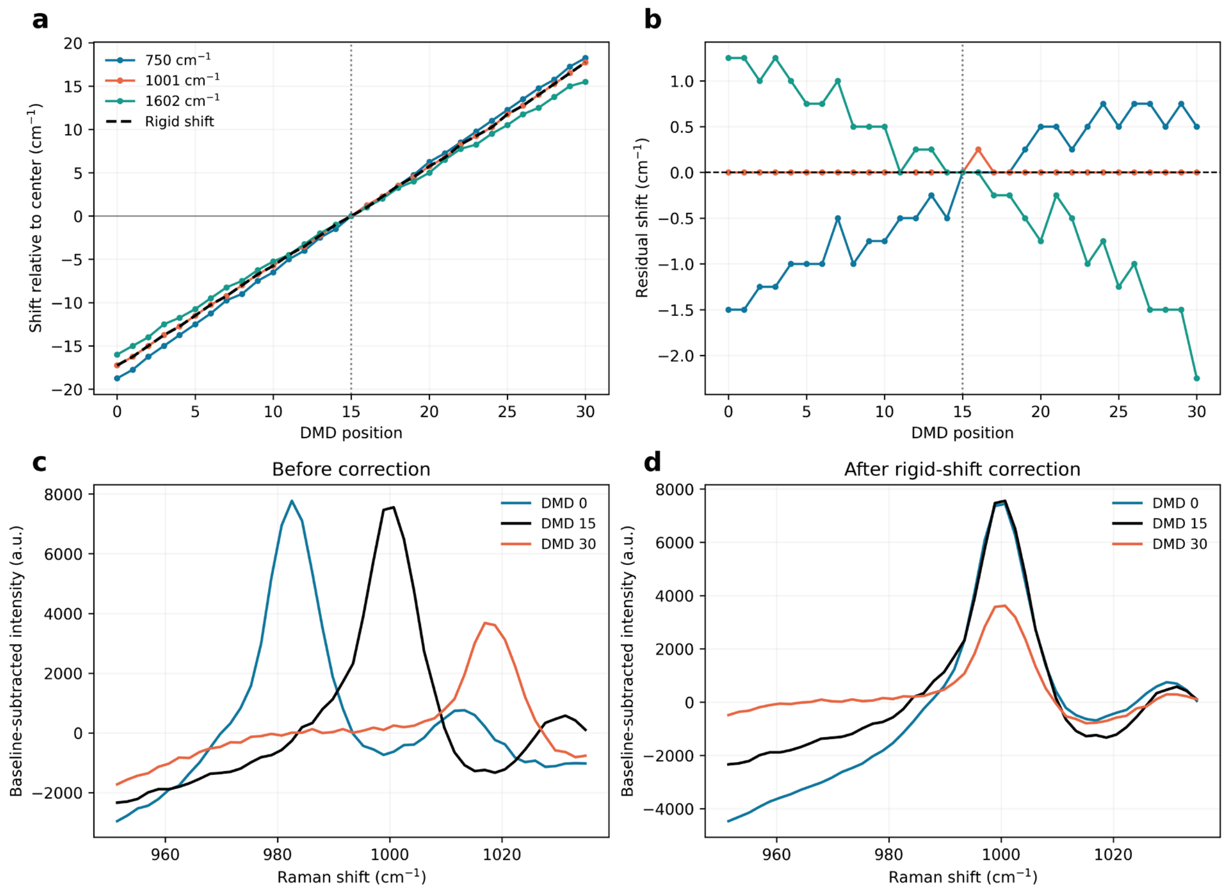


**Supplementary Figure 2 | Experimental validation of rigid spectral translation in coded-aperture Raman detection.** (a) Position-dependent spectral displacement measured across 31 DMD positions spanning the physical encoding region, using the central position (DMD 15) as the reference. Spectral shifts were independently estimated through normalized cross-correlation in three regions centered near 750, 1001 and 1602 $cm^{-1}$, following subtraction of slowly varying spectral background contributions. The dashed black line represents the common rigid displacement, defined as the median of the three independently estimated shifts at each DMD position. The common displacement exhibited a highly linear dependence on DMD position, with a slope of 1.159 $cm^{-1}$ per calibration position and a coefficient of determination of $R^2$ = 0.99985.

(b) Residual spectral displacements calculated by subtracting the common rigid shift from each independently estimated spectral displacement. The three investigated spectral regions showed closely consistent behavior, with a median interband spread of 1.50 $cm^{-1}$ and a maximum of 2.75 $cm^{-1}$ across all 31 positions. Although small systematic differences between spectral regions were observed, their magnitude remained substantially below the experimentally measured coded-aperture spectral linewidth of approximately 10 $cm^{-1}$ (Supplementary Fig. S3). These observations support a rigid translation as an effective first-order representation of the spatial–spectral coupling across the investigated Raman range.

(c,d) Representative spectral features near 1001 $cm^{-1}$ from DMD positions 0, 15 and 30, shown before (c) and after (d) application of the common rigid spectral correction. The initially displaced features converge toward the same Raman-shift position following correction, demonstrating that the dominant position-dependent spectral displacement can be compensated through a single translation. The correction preserves the relative spectral intensities without requiring intensity normalization.

The 31 calibration positions cover the physical encoding region, which is sampled at finer spatial intervals in the 63-element Hadamard configuration. Importantly, the measured displacement curves were used to validate the rigid-shift approximation rather than to provide a fixed correction map for the experimental reconstruction. Instead, spectral registration in TIRR is performed using the intrinsic Raman band of the sapphire substrate near 650 $cm^{-1}$, which provides an internal reference within the reconstructed spectra. This strategy allows the correction to be determined directly from the acquired Raman data, avoiding the need for a separate position-dependent calibration for each optical alignment. Because the correction is referenced to a spectral feature of the substrate itself, it can potentially accommodate moderate changes in optical alignment without requiring recalibration of the complete spatial–spectral mapping. The combination of experimentally validated near-rigid displacement and internally referenced spectral registration therefore provides a straightforward and flexible approach to correcting the spatial–spectral coupling introduced by the coded-aperture detection geometry.

### Supplementary Figure 3

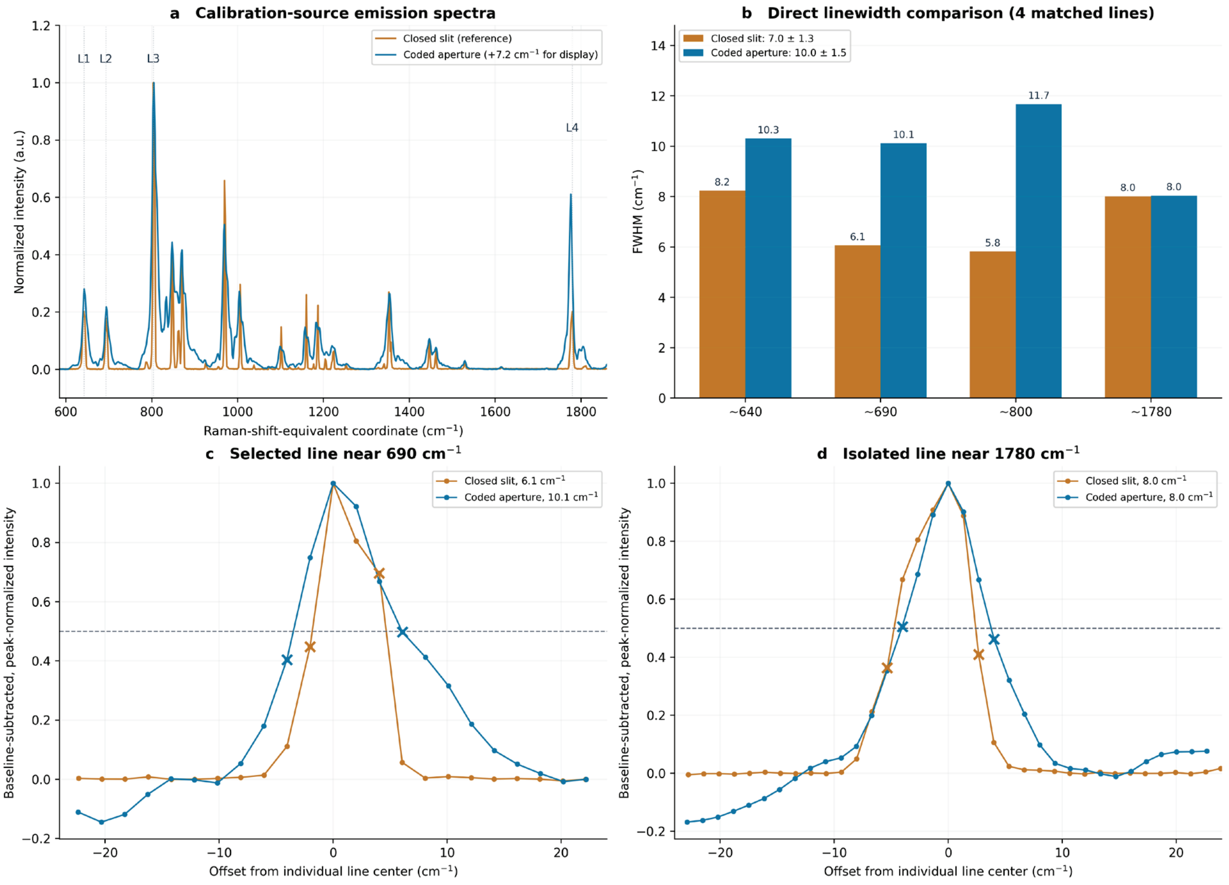


Experimental spectral-resolution characterization of the TIRR detection system. (a) Calibration-source emission spectra acquired with the spectrograph slit closed (orange) and through the coded-aperture configuration (blue). Four selected emission features (L1–L4), located near 640, 690, 800, and 1780 $cm^{-1}$, are indicated. (b) Comparison of the full widths at half maximum (FWHM) measured directly from the detector samples for each selected feature. The mean FWHM was 7.03 ± 1.26 $cm^{-1}$ with the slit closed and 10.04 ± 1.50 $cm^{-1}$ with coded-aperture detection (mean ± standard deviation across four lines). (c,d) Representative baseline-corrected and peak-normalized emission profiles near 690 and 1780 $cm^{-1}$, respectively. The individual peaks were centered for visual comparison. Horizontal dashed lines indicate 50% of the peak height, and crosses mark the measured detector samples used to calculate the FWHM. No line-shape fitting or spectral smoothing was applied. All linewidths were measured before spectral registration.

### Supplementary Figure 4

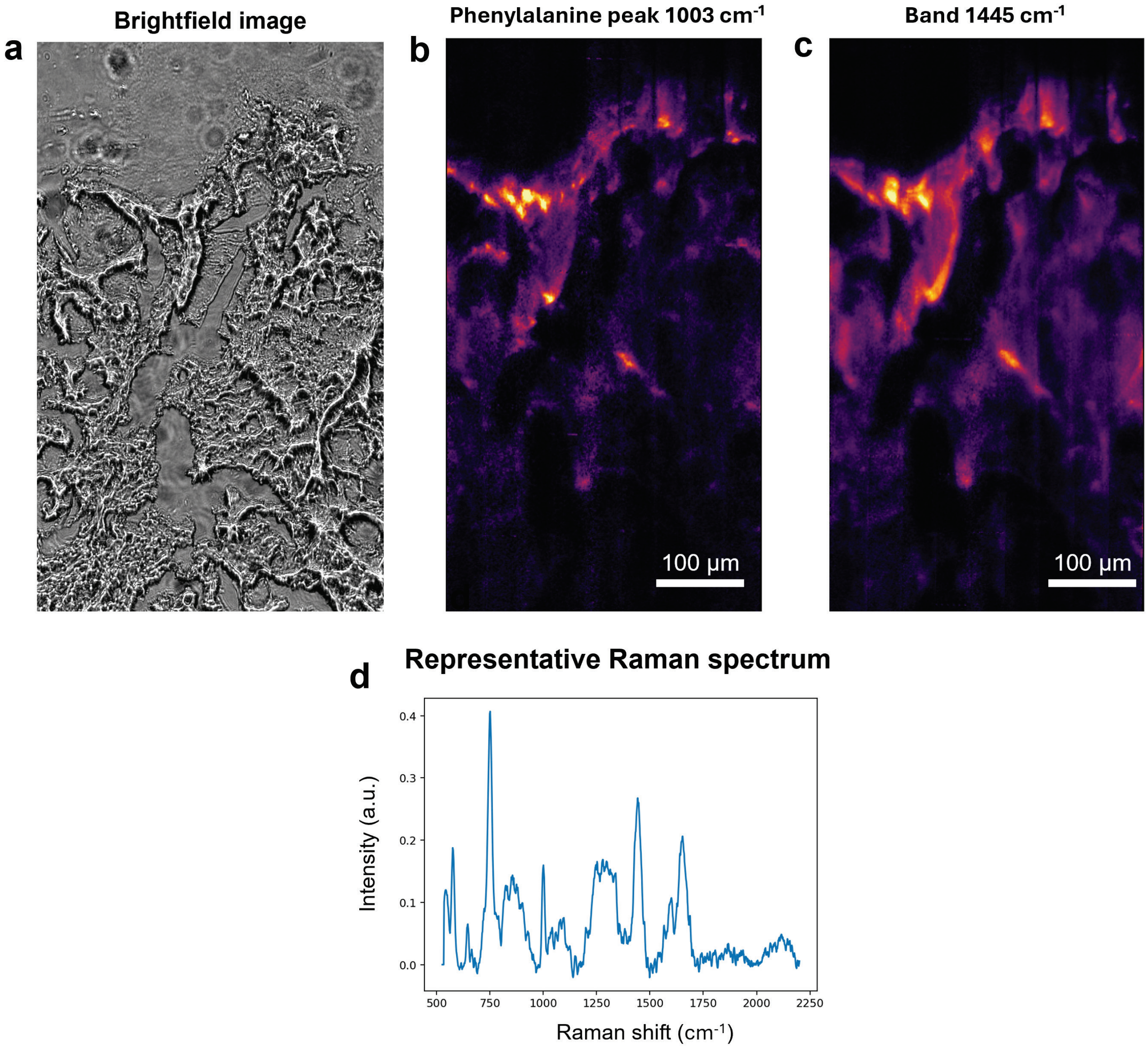


**Supplementary Figure 4 | Additional TIRR imaging example in human colon tissue.** (a), Brightfield image of an additional human colon tissue section acquired on the sapphire substrate before TIRR measurement. (b,c), TIRR Raman intensity maps extracted from the reconstructed hyperspectral cube at approximately 1003 $cm^{-1}$ and 1445 $cm^{-1}$, respectively, showing spatially heterogeneous vibrational contrast across the tissue. The map at 1003 $cm^{-1}$ highlights variations associated with the phenylalanine band, whereas the 1445 $cm^{-1}$ map emphasizes broader $CH_2$/$CH_3$ vibrational contributions. (d), Representative Raman spectrum extracted from the same dataset, illustrating recovery of the full spontaneous Raman fingerprint rather than selected discrete bands alone. This additional example supports the reproducibility of TIRR hyperspectral imaging across independent colon tissue specimens.

**Supplementary Figure 5**

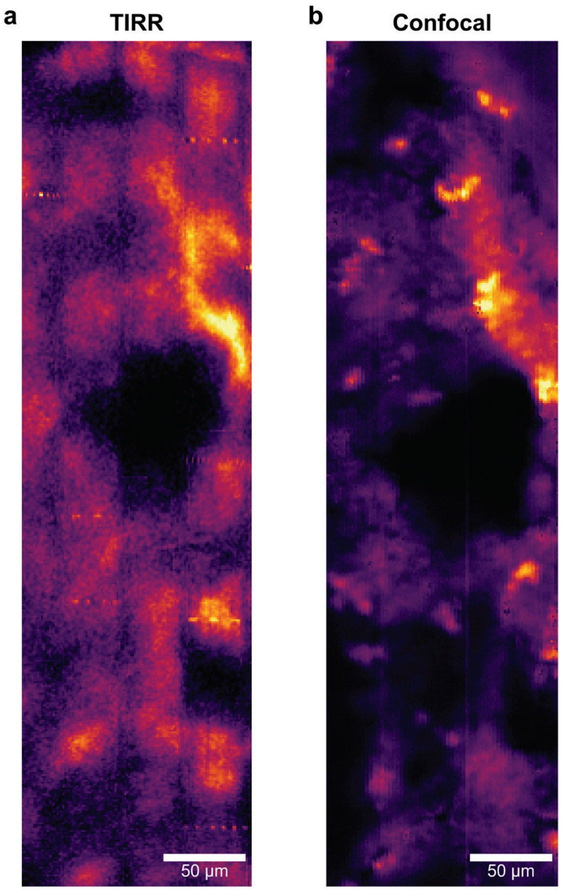


**Supplementary Figure 5 | Comparison of Raman tissue images acquired using TIRR and confocal microscopy.** Raman intensity maps reconstructed near 1450 $cm^{-1}$ from the same human colon tissue specimen using (a) total internal reflection Raman (TIRR) microscopy and (b) conventional confocal Raman microscopy. Images were obtained from corresponding tissue regions and displayed at approximately matched physical scales, with independent intensity normalization to account for differences in acquisition geometry and signal magnitude. Both modalities reveal major tissue structures, including an extended region of reduced Raman intensity and adjacent localized areas of enhanced contrast. Differences in the visibility of finer morphological features reflect the distinct spatial sampling and imaging characteristics of the two techniques. The displayed images were generated by averaging the spectral intensity over 20 detector channels surrounding the 1450 $cm^{-1}$ band, predominantly associated with $CH_2/CH_3$ deformation vibrations of proteins and lipids. The images provide a morphological reference for the spectral acquisition-efficiency benchmark presented in Fig. 4; spectral SNR was evaluated separately using the phenylalanine band near 1003 $cm^{-1}$. Scale bars, 50 µm.

### Supplementary Figure 6

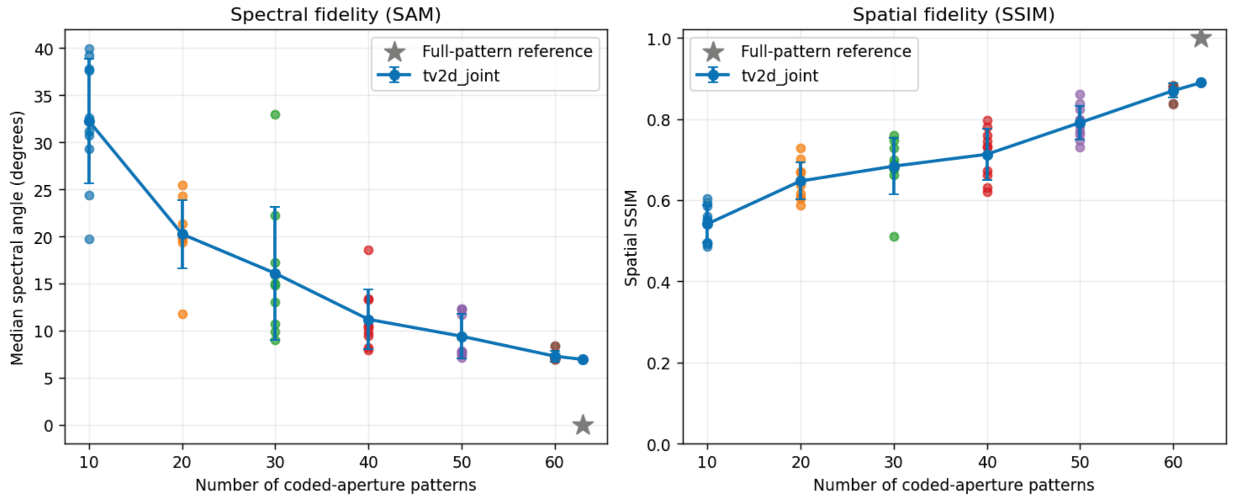


Influence of the number of Hadamard measurements on compressed TIRR reconstruction fidelity. (a) Spectral reconstruction fidelity quantified using the median pixel-wise spectral angle mapper (SAM, degrees) as a function of the number of coded-aperture patterns. (b) Spatial reconstruction fidelity evaluated using the structural similarity index (SSIM) calculated from spectral-norm intensity maps. For each measurement budget of 10, 20, 30, 40, 50 and 60 patterns, ten randomly selected subsets of the complete 63-pattern encoding matrix were independently reconstructed using joint spatial–spectral total variation regularization. Individual points represent the reconstruction metrics obtained from each subset; solid blue lines and error bars indicate the mean ± standard deviation across the ten trials. The 63-pattern TV reconstruction is also shown. Grey stars indicate the expected values when comparing the conventional full-pattern reference against itself (SAM = 0°; SSIM = 1). Increasing the measurement budget progressively improves spectral and spatial reconstruction fidelity, while variations among randomly selected subsets reveal the influence of mask composition, particularly at intermediate sampling levels.

**Supplementary Figure 7**

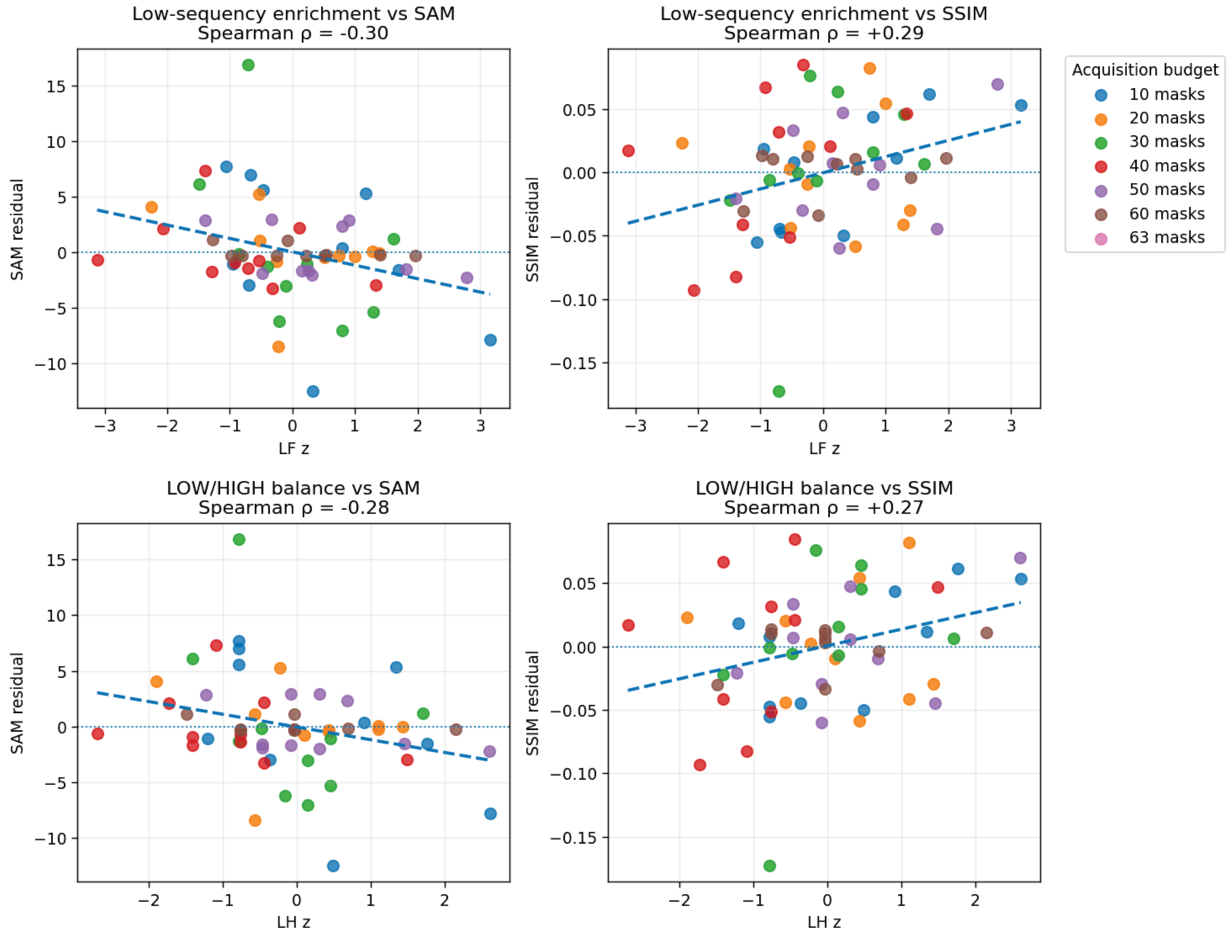


Association between Hadamard mask composition and compressed reconstruction fidelity. (a,b) Relationship between standardized low-sequency (LF) enrichment and reconstruction fidelity, expressed as residual median spectral angle mapper (SAM; a) and residual structural similarity index (SSIM; b). (c,d) Corresponding relationships for the standardized LOW/HIGH sequency balance, defined by the relative representation of low- and high-sequency masks. Each point represents one randomly selected Hadamard subset, with colors indicating the number of acquired patterns. LF enrichment quantifies the reduction in mean sequency relative to random selection, normalized by its finite-population sampling variability. SAM and SSIM residuals were obtained by subtracting the mean metric within each measurement budget to remove the dominant dependence on pattern number. Spearman rank correlation coefficients are reported for each relationship: LF enrichment versus SAM (ρ = −0.30) and SSIM (ρ = +0.29); LOW/HIGH balance versus SAM (ρ = −0.28) and SSIM (ρ = +0.27). Dashed lines indicate linear trends for visualization, and horizontal dotted lines indicate zero residual. Positive LF enrichment is associated with improved spectral and spatial reconstruction fidelity. Full 63-pattern subsets were excluded from this analysis because mask composition has no sampling variability when all patterns are included.

### Supplementary Figure 8

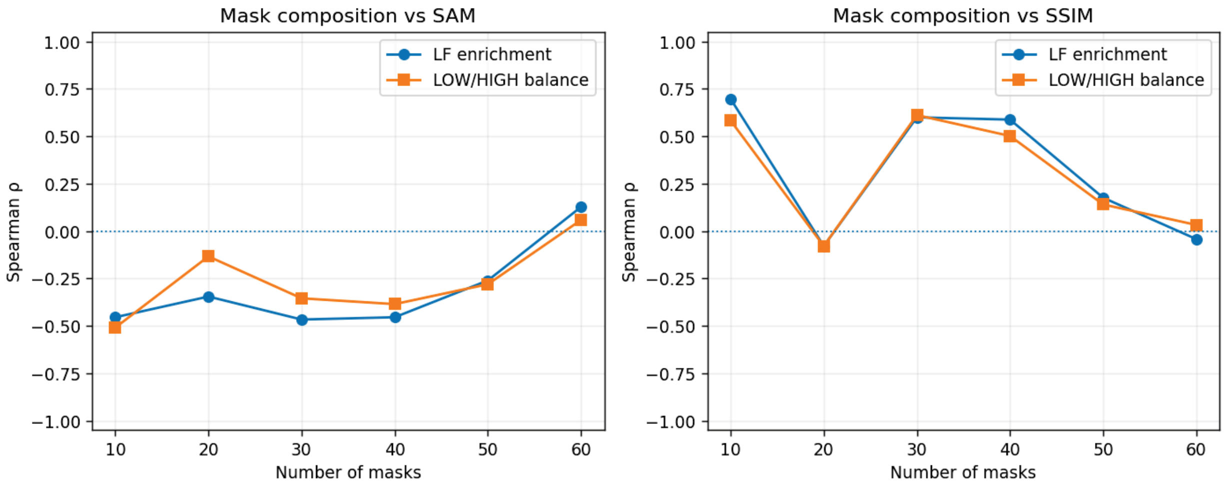


Dependence of mask-composition correlations on the number of acquired Hadamard patterns. Spearman rank correlation coefficients between two mask-composition descriptors and (a) spectral fidelity, quantified using the median spectral angle mapper (SAM), and (b) spatial fidelity, quantified using the structural similarity index (SSIM), as a function of the measurement budget. Blue circles represent standardized low-sequency (LF) enrichment, whereas orange squares represent the LOW/HIGH balance. At each measurement budget, correlation coefficients were calculated across ten independently selected random mask subsets. Negative correlations with SAM indicate an association between enrichment in low-sequency patterns and reduced spectral error, whereas positive correlations with SSIM indicate improved spatial similarity. Both descriptors exhibit consistent associations with improved reconstruction fidelity at 30 and 40 patterns, with weaker or less consistent relationships at other sampling levels. The horizontal dotted line indicates zero correlation. The complete 63-pattern condition is omitted because all masks are necessarily included and mask composition cannot vary. These correlations are exploratory and describe subset-dependent reconstruction behavior rather than independent biological replication.

**Supplementary Table 1 | Spectral and spatial reconstruction fidelity across different Hadamard sampling budgets.**

| Patterns | SAM mean ± SD (°) | SAM median [range] (°) | SSIM mean ± SD | SSIM median [range] |
|---|---|---|---|---|
| 10 | 32.23 ± 6.65 | 31.87 [19.72–39.94] | 0.541 ± 0.046 | 0.551 [0.486–0.603] |
| 20 | 20.24 ± 3.62 | 20.08 [11.78–25.49] | 0.647 ± 0.046 | 0.644 [0.588–0.729] |
| 30 | 16.08 ± 7.08 | 14.89 [9.01–32.92] | 0.684 ± 0.069 | 0.687 [0.511–0.760] |
| 40 | 11.20 ± 3.15 | 10.38 [7.95–18.55] | 0.713 ± 0.063 | 0.732 [0.620–0.798] |
| 50 | 9.41 ± 2.38 | 7.81 [7.15–12.35] | 0.791 ± 0.041 | 0.789 [0.731–0.861] |
| 60 | 7.30 ± 0.58 | 7.02 [6.99–8.40] | 0.870 ± 0.018 | 0.878 [0.836–0.883] |
| 63 | 6.95 ± 0.00 | 6.95 [6.95–6.95] | 0.889 ± 0.000 | 0.889 [0.889–0.889] |

Summary of reconstruction performance obtained using joint spatial–spectral total variation regularization and randomly selected subsets of the 63-pattern Hadamard encoding matrix. For each sampling budget, ten trials were evaluated against the conventional full-pattern reconstruction using the median pixel-wise spectral angle mapper (SAM) and structural similarity index (SSIM) of spectral-norm intensity maps.

For 10–60 patterns, statistics were calculated from ten randomly selected subsets per measurement budget. At 63 patterns, the ten recorded trials contain the same complete mask set and yield identical reconstruction metrics; they do not constitute ten distinct mask selections. Lower SAM and higher SSIM indicate better reconstruction fidelity.

For comparison, the deterministic 30-pattern low-sequency subset achieved SAM = 7.37° and SSIM = 0.8633 in the colon development dataset, and SAM = 7.24° and SSIM = 0.8846 in the independent skin dataset (Fig. 5). These deterministic reconstructions are reported separately from the random-trial statistics.

**Supplementary Table 2 | Quantitative comparison of senescence-associated TIRR Raman bands in human skin.**

Cell-associated Raman band intensities were compared between p21-positive (p21+, n = 22) and p21-negative (p21−, n = 965) regions within a single human skin tissue section. Spectra were normalized using their L1 norm over the 600–1800 $cm^{-1}$ fingerprint region. Five RamanOmics-informed vibrational bands were evaluated using two-sided Mann–Whitney U tests, with p-values adjusted for multiple comparisons using the Holm procedure.

| Raman band ($cm^{-1}$) | Assignment | p21+ median | p21− median | RBC | p-value | Holm-adjusted p |
|---|---|---|---|---|---|---|
| 1134 | Lipid | 2.289 | 1.101 | +0.403 | 0.001218 | 0.003048 |
| 940 | Collagen | 15.001 | 18.640 | −0.461 | 0.000216 | 0.000863 |
| 1163 | Collagen | 2.579 | 3.837 | −0.490 | 0.000084 | 0.000421 |
| 1440 | Lipid | 33.459 | 30.978 | +0.269 | 0.030727 | 0.030727 |
| 1393 | Amino acid | 12.293 | 16.005 | −0.409 | 0.001016 | 0.003048 |

Median intensities are expressed in units of $\times 10^{-4}$ L1-normalized Raman intensity. Rank-biserial correlation (RBC) quantifies the direction and magnitude of the group separation, with positive values indicating higher intensities in p21+ regions and negative values indicating lower intensities. Holm adjustment was applied across the five investigated bands. All eligible cell-associated measurements were included in the statistical analysis. Because the data originate from one tissue section of a single donor, the statistical comparisons are exploratory at the cellular level and do not establish biological reproducibility across donors.

**Supplementary Table 3 | Oligonucleotide sequences for STARmap-ISH targeting CDKN1A (p21) and CDKN2A (p16).**

| Target gene | Oligonucleotide | Sequence (5'–3') | 5' modification |
|---|---|---|---|
| CDKN1A | Primer 1 | GGCCTTGCTGCCGCATGGGTTTAATGTTATCTT | — |
| CDKN1A | Padlock 1 | ACATTATGACGGACATCCCCAGCCGGTTCAATTATC GACGAAACTTCGTAATTAAGATA | 5' phosphate |
| CDKN1A | Primer 2 | GTCTTCCTCTGCTGTCCCCTAATGTTATCTT | — |
| CDKN1A | Padlock 2 | ACATTACAGCAGAGCAGGTGAGGTGAATTATCG ACGAAACTTCGTAATTAAGATA | 5' phosphate |
| CDKN1A | Primer 3 | TTCCTCTTGGAGAAGATCAGCCTAATGTTATCTT | — |
| CDKN1A | Padlock 3 | ACATTAGCGTTTGGAGTGGTAGAAATCTAATTATCGA CGAAACTTCGTAATTAAGATA | 5' phosphate |
| CDKN1A | Decoding probe | CGACGAAACTTCGTAATT | 5' Alexa Fluor 488 |
| CDKN2A | Primer 1 | TCATGACCTGGATCGGCCTTAATGTTATCTT | — |
| CDKN2A | Padlock 1 | ACATTACGACCGTAACTATTCGGTGCGTAATTATa atctcctctgggtaaaaAAGATA | 5' phosphate |
| CDKN2A | Primer 2 | TCCGCGGCATCTATGCGGGTAATGTTATCTT | — |
| CDKN2A | Padlock 2 | ACATTATGGTTACTGCCTCTGGTGCCAATTATaatctcc tctgggtaaaaAAGATA | 5' phosphate |
| CDKN2A | Primer 3 | GCAGGTACCGTGCGACATCGCGTAATGTTATCTT | — |
| CDKN2A | Padlock 3 | ACATTATGGCCCAGCTCCTCAGCCAGAATTATaatc tcctctgggtaaaaAAGATA | 5' phosphate |
| CDKN2A | Decoding probe | aatctcctctgggtaaaa | 5' Alexa Fluor 647 |

Sequences are given in the 5'–3' orientation, without vendor-specific modification codes. The 5'-end chemical modifications are indicated separately. Numbered primers and padlocks form corresponding probe pairs for each gene. Nucleotide capitalization has been preserved from the supplied oligonucleotide list. CDKN1A (p21) annotations were used for the Raman–senescence analysis shown in Fig. 6; CDKN2A (p16) probes are included for completeness.
